# Resolution of glycogen and glycogen-degrading activities reveals correlates of *Lactobacillus crispatus* dominance in a cohort of young African women

**DOI:** 10.1101/2022.03.29.486257

**Authors:** Karen V. Lithgow, Athena Cochinamogulos, Kevin Muirhead, Shaelen Konschuh, Lynda Oluoch, Nelly R. Mugo, Alison C. Roxby, Laura K. Sycuro

**Affiliations:** Department of Microbiology, Immunology and Infectious Diseases, University of Calgary, Calgary CANADA; Department of Biochemistry and Molecular Biology, University of Calgary, Calgary CANADA; Kenya Medical Research Institute, Nairobi KENYA; Department of Global Health, University of Washington, Seattle USA; Department of Medicine, University of Washington, Seattle USA; Department of Epidemiology, University of Washington, Seattle WA USA; Vaccine and Infectious Disease Division, Fred Hutchinson Cancer Research Center, Seattle WA USA; Calvin, Phoebe & Joan Snyder Institute for Chronic Diseases, Cumming School of Medicine, University of Calgary, Calgary CANADA; Alberta Children’s Hospital Research Institute, Cumming School of Medicine, University of Calgary, Calgary CANADA; International Microbiome Centre, University of Calgary, Calgary CANADA

**Keywords:** vaginal microbiome, *Lactobacillus* dominance, *Lactobacillus crispatus*, glycogen, amylase, pullulanase, lactic acid, African women, adolescence

## Abstract

**Background:** A healthy vaginal microbiome is dominated by *Lactobacillus* species that produce lactic acid, lowering vaginal pH and limiting colonization by pathogens. *Lactobacillus* dominance (LD) is established during puberty, but many women, especially those of Black race, lose LD during their reproductive years. Glycogen is thought to be a key host nutrient that supports vaginal lactobacilli and their fermentative lactic acid production, but mechanisms of glycogen utilization by *Lactobacillus* species are incompletely understood. By partitioning glycogen and glycogen-derived maltodextrin, as well as the activity of glycogen-degrading pullulanase enzymes, this work refines understanding of vaginal glycogen catabolism and identifies correlates of LD.

**Results:** Vaginal swab samples were collected from a cohort of young women with limited sexual experience in Thika, Kenya (N=17, ages 17–20). Metagenomic profiling of the vaginal microbiome revealed that most samples exhibited LD, particularly dominant *Lactobacillus crispatus*. Amylopullulanase activity, cleavage of glycogen α-1,4 and α-1,6 linkages by individual/multifunctional enzymes, showed a significant positive correlation with glycogen-derived maltodextrin, but no relationship with *L. crispatus* dominance. Pullulanase activity, which specifically targets glycogen α-1,6 linkages, was 3-fold higher in *L. crispatus-*dominated samples and significantly correlated with D-lactic acid levels. Metagenomics and targeted PCR revealed that 36% of *L. crispatus*-dominated metagenomes from our African cohort lacked a functional *L. crispatus* pullulanase (*pulA*) gene, a 3-fold higher frequency of gene loss than that seen in metagenomes from European and North American women. Our findings suggest *pulA* gene loss or inactivation may correspond with reductions in *L. crispatus* abundance, pullulanase activity and lactic acid levels compared to samples dominated by *pulA*-competent *L. crispatus*.

**Conclusions:** Our results indicate that although amylase activity drives the accumulation of glycogen catabolites in vaginal fluid, pullulanase appears to specifically contribute to maximal D-lactic acid production by *L. crispatus*. However, this is only possible when a functional *pulA* gene is present, which was not the case in a substantial proportion of young African women with dominant *L. crispatus*. Scaling this analysis to a larger cohort will address whether genomic and enzymatic indicators of *L. crispatus* pullulanase activity are predictive of sustained LD and vaginal health.

## Background

The optimal vaginal microbiome of reproductive aged women is typically characterized by low species diversity, low pH and dominance by lactic acid-producing *Lactobacillus* species (spp.), including *Lactobacillus crispatus, Lactobacillus jensenii*, *Lactobacillus gasseri and Lactobacillus iners* [1–3]. Loss of protective *Lactobacillus* spp. contributes to a dysbiotic shift in the vaginal microbiome that can lead to bacterial vaginosis (BV), a clinical condition characterized by overgrowth of diverse anaerobic bacteria [4, 5]. Bacterial vaginosis is associated with increased risk of preterm birth [6–8], genital tract cancers [9–11] and acquisition and transmission of sexually transmitted infections (STI), including human immunodeficiency virus (HIV) [12, 13]. Previous studies have shown that the vaginal microbiome displays compositional variance by race and ethnicity [3, 14–18]. Up to 40% of African American women do not exhibit *Lactobacillus* dominance, corresponding with a prevalence of BV that is 2–3x higher than that in Caucasian women [3, 16, 17, 19]. In adult Black women with dominant *Lactobacillus* spp., the less protective species *L. iners* is most prevalent (58–65%), with *L. crispatus* and *L. jensenii* following at 20% and 13% prevalence, respectively [14, 18, 20, 21]. A high incidence of BV and low prevalence of *L. crispatus* have also been documented in African women, increasing their risk of HIV acquisition [21–23]. Furthermore, loss of *L. crispatus* and increased BV susceptibility have been linked to sexual activity in both girls and women [18, 24–27]. However, recent studies of adolescent girls from Tanzania and Kenya revealed that samples captured around the time of sexual debut exhibit a higher frequency of *L. crispatus* dominance [27–29]. This highlights a critical need to continue studying adolescent populations, in which fewer environmental exposures (sex, vaginal hygiene practices, hormonal contraceptives) confound relationships between nutritive biomolecules and *L. crispatus* dominance of the vaginal niche.

A key aspect of *Lactobacillus* dominance is the fermentative production of lactic acid by *Lactobacillus* spp., which acidifies the vaginal microenvironment and protects against genital tract colonization by pathogens or BV-associated bacteria [2, 30–37]. While the L-lactic acid isomer is produced by human cells and vaginal lactobacilli, the D-lactic acid isomer is only produced by *L. jensenii, L. gasseri, L. paragasseri* and *L. crispatus* [38–40]. Since free glycogen in vaginal fluid correlates with *Lactobacillus* dominance and low pH [41, 42], glycogen is proposed to be a key nutrient supporting vaginal lactobacilli and their lactic acid production. Glycogen is a large, highly branched D-glucose polymer that contains α-1,4 glycosidic bonds within linear chains and α-1,6 linkages at branch points [43]. Human α-amylase enzymes present in vaginal fluid can metabolize glycogen α-1,4 linkages to generate breakdown products, including maltose, maltotriose and maltodextrin (longer unbranched D-glucose polymers). Several *Lactobacillus* spp. have been shown to directly utilize these glycogen breakdown products for growth [44, 45]. Recent work by van der Veer et al. confirms that vaginal isolates of *L. crispatus* can utilize glycogen as a sole carbon source [46], in contrast with prior work that concluded *Lactobacillus* spp. do not directly utilize glycogen [44, 47–49]. Intriguingly, van der Veer et al. showed that rapid growth of *L. crispatus* on glycogen correlated with the presence of a functional α-1,6 targeting type I pullulanase (*pulA*) gene; *L. crispatus* isolates containing putative inactivating deletions in the *pulA* gene were deficient for growth on glycogen [46]. While pullulanase enzymes target the α-1,6 glycogen branched linkages, endo-acting α-amylases and exo-acting α-glucosidases target the α-1,4 linkages [43]. Genes encoding both α-1,6 and α-1,4 targeting enzymes have also been identified in other *Lactobacillus* species, including, *L. iners, L. gasseri* and *L. jensenii* [50–52], as well as BV-associated bacteria [51, 53]. Furthermore, glycogen and maltodextrin degrading enzymes from *Lactobacillus* spp. and BV-associated bacteria have been detected in human vaginal fluid proteomes [51]. However, the competitive/cooperative utilization of glycogen and maltodextrin by *Lactobacillus* spp. and BV-associated bacteria is not understood.

Although the temporal and correlative link between *Lactobacillus* dominance, glycogen and lactic acid has been described in several studies [41, 42], our work shows that commonly-employed glycogen quantification kits non-specifically detect both glycogen and glycogen-derived glucose polymers, including maltose, maltotriose and maltodextrin in vaginal samples. Similarly, most enzyme activity kits do not distinguish α-1,4 and α-1,6 targeting enzymes. In this study, we analyzed vaginal swab specimens from a cohort of 17 adolescent girls and young women (AGYW) from Thika, Kenya. The aim of this pilot analysis was to demonstrate that partitioning glycogen catabolites and enzyme activities can deepen our understanding of glycogen utilization and provide clues as to how *L. crispatus* dominance may be therapeutically sustained in the vaginal niche.

## Methods

### Kenyan cohort of adolescent girls and young women (AGYW)

Samples for this pilot cohort were collected from a prospective sexual health study of AGYW with limited sexual experience in Thika, Kenya (full study details are provided in [54]). Girls who met the following criteria and provided written informed consent with their guardian present were enrolled between 2014 and 2016: 1) age 16–19; 2) HIV and HSV-2 seronegative; 3) willing to undergo genital examination; 4) ≤1 sexual partners (total) to date. All samples used in this analysis were from the baseline clinical visit at which the participants underwent a genital exam and provided vaginal swab specimens. Demographic and sexual history data was gathered using questionnaires. Participant demographic and clinical metadata can be found in Table 1 and Table S1, Additional File 2. Study approval was obtained from the Kenya Medical Research Institute Scientific Ethics Review Unit, the University of Washington Institutional Review Board and the Conjoint Health Research Ethics Board (CHREB) at the University of Calgary.

**Table 1.** Characteristics of adolescent girls and young women (AGYW) enrolled in a prospective sexual health study in Thika, Kenya

| **Baseline Characteristic** | **Pilot Cohort<sup>a</sup>**<br>**(N = 17)** | **Full Cohort**<br>**(N = 400)** |
| --- | --- | --- |
| Age at enrollment | 19 (18–20) | 18.6 (17.6–19.4) |
| Years of school completed | 12 (10.5–12) | 12 (10–12) |
| Socioeconomic factors |  |  |
| *Running water* | 5 (29.4) | 191 (47.8) |
| *Electricity* | 6 (35.3) | 277 (69.3) |
| *Concrete floor* | 10 (58.8) | 290 (72.5) |
| Sexually active<sup>b</sup> | 4 (23.5) | 78 (19.5) |
| Age at first sex | 18 (15.8–18) | 18 (18–19) |
| Number of times sex in last 3 months | 1 (0–1) | 1 (0–2) |
|  | **Pilot Cohort**<br>**BV/STI Tested**<br>**(N = 17)** | **Full Cohort**<br>**BV/STI Tested<sup>c</sup>**<br>**(N = 375/373)** |
| BV negative; Nugent score 0–6<sup>d,e</sup> | 17 (100) | 354 (94.4) |
| STI<sup>f</sup> | 1 (5.9)<sup>g</sup> | 49 (13.1) |

**Data expressed as median (IQR) or N (%)**<sup>a</sup> A subset of the full cohort that did not return after the baseline visit.<sup>b</sup> All sexual history data were self-reported.<sup>c</sup> 375 participants in the full study were tested for bacterial vaginosis at baseline; 373 were tested for chlamydia, gonorrhea and trichomonas at baseline. Reasons for not testing include menstruation and other factors that prevented vaginal swab collection.<sup>d</sup> Bacterial vaginosis (BV) was diagnosed at Kenyatta National Hospital using Nugent scoring; scores $\geq 7$ were considered positive for BV and those $< 7$ negative for BV.<sup>e</sup> Nugent scores were previously published for the full cohort [54]. Vaginal microbiome data is, to date, only available for this pilot subset.<sup>f</sup> Sexually transmitted infections (STI) is a combined variable including chlamydia, gonorrhea and trichomoniasis. All participants were HSV and HIV seronegative at baseline.<sup>g</sup> Trichomoniasis was the only STI present in the pilot cohort at baseline.

### Biospecimen processing

After collection, samples were stored at −80°C and transported on dry ice from the study site to the University of Washington, and then on to the University of Calgary. Saline (0.9% NaCl) was prepared with UV irradiated H_2_O (HyClone; Cytiva Life Sciences, Marlborough, MA, USA) and filtered with a 100 kDa molecular weight cut-off (MWCO) Amicon Ultra-15 filter unit (Millipore, Oakville, ON, CA). Vaginal swabs were eluted in 1.5 mL sterile filtered saline by thawing at room temperature for 5–10 minutes (min), then vigorously vortexed for 1 min to release the remaining biomass; subsequent swab processing steps were conducted at 4°C. Swab eluates were centrifuged at 18,407 × g for 10 min and resulting cell-free supernatant and pellet samples stored at −80°C. Swabs were processed in batches of 9, including a sham swab as a negative control (Puritan sterile foam tipped applicator swab; Guilford, Maine, USA).

### Biomolecule analyses

#### Total protein

Total protein content in cell-free swab supernatants was measured using the Pierce^TM^ BCA protein assay according to the manufacturer’s instructions (ThermoFisher Scientific, Burnaby, BC). Briefly, 25 µL aliquots of swab supernatants or bovine serum albumin (BSA) standards diluted in 0.9% saline were added to a 96 U-well plate (Greiner Bio-One, Monroe, NC, USA) in technical duplicate. Working reagent (200 µL/well) was added prior to incubation at 37°C for 30 min. The plate was then cooled to room temperature before the optical density at 562 nm (OD562) was measured in a BioTek Synergy H1 plate reader (BioTek Instruments, Inc, Winooski, VT) after a 5 second orbital shake. To account for differences in swab load/biomass that can result from sampling variability, all biomolecule measurements (lactic acid, glycogen/maltodextrin and enzyme activities) were normalized to the total protein concentration of each vaginal swab supernatant (Tables S2–S3, Additional File 2).

#### Lactic acid

D- and L-lactic acid were quantified in cell-free swab supernatants using the sequential method of the D-/L-Lactic Acid Rapid Assay Kit (Megazyme, Darmstadt, Germany). Adapted from the manufacturer’s instructions for a 96 well plate format, 50 µL of D-/L-lactic acid standards and swab supernatants were aliquoted into a 96 U-well plate (Greiner Bio-One) in technical duplicate. Samples and standards were mixed with D-glutamate-pyruvate transaminase (2 µL/well), NAD+ (9 µL/well), buffer (45 µL/well) and water (83 µL/well), and the background absorbance (OD340) was measured in a BioTek Synergy H1 plate reader. D-lactic acid was quantified by adding D-lactate dehydrogenase (5 µL) to each well and measuring OD340 after a 20 min incubation following a 5 second orbital shake. Next, 5 µL of L-lactate dehydrogenase was added to each well of the same plate and the same steps were followed for quantification of L-lactic acid. Total lactic acid was considered the sum of D- and L-lactic acid.

#### Glycogen and maltodextrin

The specificity of a fluorometric glycogen assay kit (BioVision, Milpitas, CA) was evaluated by testing 3.2 µg/mL solutions of glycogen (BioVision), maltodextrin (Sigma-Aldrich), maltotriose (Sigma-Aldrich, St. Louis, MO) and maltose (ThermoFisher Scientific) (Figure S1, Additional File 1). These solutions, alone and in combination, were then used to validate that size exclusion improves the kit’s specificity for glycogen. Carbohydrate solutions (50 µL) were diluted in 250 µL of HyPure H_2_O (Cytiva Life Sciences, Marlborough, MA) and added to a 100 kDa MWCO Amicon Ultra-0.5 centrifugal filter unit (Millipore Sigma, Oakville, ON). Glycogen (concentrate) and maltodextrin/glucose (filtrate) were separated by centrifuging at 14,000 x g for 2.5 min at room temperature. After centrifugation, the volume of each concentrate/filtrate was adjusted to 300 µL with HyPure H_2_O. Filtrate and diluted concentrate (1:100) were added to a black optical bottom 96 well plate (10 µL/well; Greiner Bio-One) in technical duplicate and each fraction was processed separately in the presence or absence of hydrolysis enzymes according to the manufacturer’s instructions. Fluorescence (excitation/emission at 535/587 nm) was measured using a BioTek Synergy H1 plate reader. Background glucose (measured by not adding hydrolysis enzymes to the concentrate and filtrate) was subtracted from the corresponding hydrolyzed sample to measure glycogen and maltodextrin, respectively. Vaginal swab supernatants were processed using the same procedure.

#### Glycogen degrading enzyme activity

Enzyme activity in vaginal swab supernatants and cell pellets was quantified using the α-Glucosidase Activity Assay Kit (Abcam, Waltham, MA, USA), Enzchek Ultra Amylase Kit (Invitrogen, Carlsbad, CA, USA), the α-amylase Ceralpha assay kit (Megazyme, Wicklow, Ireland) and the pullulanase PullG6 Kit (Megazyme) per the manufacturers’ instructions. The specificity of the Enzchek Ultra Amylase kit was tested using control enzymes from the α-Glucosidase Activity Assay Kit (Abcam), an αmylase kit (Abcam) and the pullulanase assay kit (Megazyme). The pullulanase (Megazyme) kit was adapted for a 96 well plate. Briefly, pullulanase control enzyme dilutions (*Bacillus licheniformis* pullulanase) were used to generate a standard curve according to manufacturer instructions. These, along with swab supernatants were incubated in 1.5 mL tubes for 10 min at 40°C. In parallel, the PullG6 substrate (50 µL/well) was incubated in a 96 U-well plate (Greiner Bio-One) that was sealed to prevent evaporation. The enzyme standards (50 µL/well) and swab supernatant samples (75 µL/well) were added to the PullG6 substrate plate in duplicate wells and incubated for 30 min at 40°C. Stopping reagent (2% weight/volume Tris buffer pH 9.0) was added to each well (100 µL/well) and OD400 was measured in a BioTek Synergy H1 plate reader. Cell-associated enzyme activity was evaluated in swab pellets that had been resuspended in 50 µL of 0.9% NaCl; samples 2, 7, 13 and 17 did not have adequate sample remaining and were excluded. Reported measurements account for the volume of swab eluate used to generate the pellet, allowing for direct comparison to the activity in swab supernatants.

#### Human α-amylase concentration

The concentration of host α-amylase in swab supernatants was measured using a pancreatic α-amylase sandwich ELISA (Abcam, Toronto, ON, CA). The α-amylase standard (50 µL) and swab supernatants (75 µL) were added to the provided microplate in technical duplicate and the assay performed according to manufacturer’s instructions. A BioTek Synergy H1 plate reader was used for colorimetric quantification (OD450) after a 5 second orbital shake.

### DNA extraction and quantitative polymerase chain reaction (qPCR)

Genomic DNA was extracted from vaginal cell pellets using the DNeasy PowerSoil Pro QIAcube HT Kit on a QIAcube HT instrument (Qiagen, Germantown, MD). Sham swabs were included to monitor for contamination of swab prep and extraction materials. All extracts were assessed for the presence of PCR inhibitors by spiking in and quantitatively amplifying a synthetic jellyfish aequorin gene fragment [55]. Total bacterial load was assessed using broad range 16S rRNA gene qPCR with the following primers/probes: 16SBR_337F, 5’-ACTCCTRCGGGAGGCAGCAG-3’; 16SBR_793R, 5’-GGACTACCVGGGTATCTAATCCTGTT-3’; 16SBR_511R_FAM/TAM, 5’-/56-FAM/CGTATKACCGCGGCTGCTGGCAC/36-TAMSp/-3’ (procured as PrimeTime qPCR Probe assays with 2:1 primers:probe ratio, Integrated DNA Technologies, Coralville, IA). An *Escherichia coli* 16S rRNA gene fragment was cloned using a TOPO™ TA Cloning Kit (ThermoFisher Scientific) and the resulting plasmid used to generate the standard curve. Standard curve and extract reactions were set up in technical duplicate using TaqMan™ Environmental Master Mix 2.0 (ThermoFisher Scientific) and run on a StepOne Plus Real-Time PCR System (Applied Biosystems, ThermoFisher Scientific) with the following parameters cycled 40 times: 15 second denature at 95°C, 1 min anneal/extend at 60°C. 16S rRNA gene copies detected in a given volume of DNA extraction were volumetrically converted to copies/mL swab eluate and then to copies/swab.

### Shotgun metagenomic sequencing

Genomic DNA from vaginal cell pellets and the ZymoBIOMICS Microbial Community Standard (Zymo Research, Irvine, CA) was cleaned and concentrated with either sparQ PureMag beads (QuantaBio, Beverly, MA) or the Genomic DNA Clean & Concentrator-10 Kit (Zymo Research, Irvine, CA), according to the manufacturer’s instructions. Shotgun libraries were prepared from 50 ng genomic DNA using the QIAseq FX DNA Library Kit (Qiagen, Germantown, MD), according to manufacturer’s instructions, with fragmentation times of 10–15 min, 6–8 cycles of PCR amplification and sample-specific linkage to combinatorial dual indices. SparQ beads were used for all bead clean-up steps and to fine-tune library size distribution with peak fragment sizes of 300–500 base pairs (bp). Libraries were pooled at 3 nM and sequenced on a HiSeq X instrument (Illumina, San Diego, CA) to generate 150 bp paired-end reads (Psomagen, Rockville, MD).

### Metagenomic data processing and taxonomy classification

Raw Illumina reads were subjected to adapter trimming, quality filtering and human sequence decontamination using a custom, publicly-available snakemake pipeline (metqc: https://github.com/SycuroLab/metqc). In brief, adapters were removed using Cutadapt version 1.16 [56] and 3’/5’ end trimming and quality filtering were performed with PRINSEQ version 0.20.4 [57]. Clean reads were filtered to remove human DNA contamination using Best Match Tagger (bmtagger) version 3.101 with version 19 of the human reference genome [58]. Microbial taxonomic profiling was performed with the clean non-human reads using MetaPhlAn version 3.0.1 [59]. Bacterial species exhibiting a relative abundance ≥0.1% and prevalence ≥10% were retained in the relative abundance plot created using the ggplot2 version 3.3.3 library package [60] in R version 4.0.5 [61]. Hierarchical clustering was applied to a Bray-Curtis dissimilarity matrix calculated on MetaPhlAn3 read count data using the vegdist(bray) distance function, with specification of the UPGMA method using the hclust(average) R function, in vegan version 2.5-7 [62]. Participant metadata was incorporated into the relative abundance plot in R using pheatmap version 1.0.12 [63] with colour palettes generated using the RColorBrewer version 1.1-2 [64]. Mathematical analyses of bacterial abundance were performed on a read count table that had been adjusted for barcode hopping as follows. First the relative abundance of highly abundant/prevalent taxa unexpectedly found in the Zymo control sample due to low level barcode hopping was calculated (*L. crispatus* and *L. iners*, each representing <0.005% of the control sample relative abundance). A number of reads representing the equivalent relative abundance was then subtracted from each sample containing those two taxa; this approach scales the adjustment to the total number of reads in each sample. The net effect is that the total read count for each of these species is slightly reduced, bringing some samples that had very low read counts for these two species to 0 (undetected). Unadjusted and adjusted count tables are provided in Tables S4–S5, Additional File 2.

### Metagenome-assembled genome binning

Metagenome-assembled genome (MAG) binning was accomplished using the metaWRAP pipeline version 1.3 [65]. Metagenomic assemblies were generated for each sample with SPAdes version 3.13.0 [66] and evaluated for quality using QUAST version 5.0.2 [67]. Metagenome binning and refinement was performed using MetaBAT2 version 2.12 CheckM version 1.1.3 [71] quality assessment reported ≥50% completeness and ≤10% contamination. MAGs were annotated using prokka version 1.13 [72] and classified to the species level using the genome taxonomy database toolkit (GTDB-tk) version 1.5.0 [73].

### Presence and sequence heterogeneity of *L. crispatus pulA*

To assess the presence and allelic content of *L. crispatus* type I pullulanase (*pulA*) genes from women around the globe, whole genome assemblies from urogenital *L. crispatus* isolates were selected from the NCBI unfiltered prokaryotes list (downloaded August, 2021) by parsing the GenBank metadata for “*Homo sapiens*” and manually investigating the isolation source and referenced publications. A total of 86 *L. crispatus* urogenital tract isolate genomes were identified and verified by CheckM to be ≥90% complete and ≤10% contaminated. We also downloaded 17 publicly available *L. crispatus* MAGs binned by Pasolli et al. from human vaginal metagenomes [74–76] and annotated them as described above for Thika MAGs; only MAGs meeting our minimum quality thresholds of ≥50% completeness and ≤10% contamination were considered. Type I pullulanase genes were assessed in each isolate genome and MAG using the BLASTn program of BLAST+ version 2.9.0 [77, 78]. The full-length and functionally characterized *pulA* from *L. crispatus* isolate genome RL03 (NCBI accession NZ_NKLQ01000285) [46] was used as the query, and the resulting hits were filtered to include those with ≥95% sequence identity. *pulA* genes with <100% sequence identity or <100% query coverage were subject to further analysis using SnapGene® version 5.3.0 (GSL Biotech LLC, San Diego, CA) and InterProScan (EMBL-EBI, Cambridgeshire, UK) to evaluate potential functional consequences of any observed insertions or deletions (i.e. frameshift mutations or domain/signal peptide loss) or single nucleotide polymorphisms (SNPs, i.e. those that result in nonsense mutations). We assessed *L. crispatus* genome coverage and that of the *pulA* locus in each *L. crispatus-*containing Thika metagenome by mapping clean non-human reads from each sample to the *L. crispatus* FDAARGOS_743 reference genome (NCBI accession GCF_009730275.1) [79] using bowtie2 version 2.3.5 [80] with default parameters. Depth of coverage was calculated using samtools version 1.11 [81] and the *pulA* coding sequence region specified as positions 1986082–1989861 on the closed chromosome of the *L. crispatus* FDAARGOS_743 reference genome (NCBI accession NZ_CP046311.1). Coverage plots were generated in R using ggplot2 version 3.3.3 [60]. Finally, a targeted PCR assay was used to confirm *L. crispatus pulA* gene presence in the Thika samples. Using 20 ng of DNA extracted from the vaginal cell pellets, a near full-length *pulA* gene fragment was amplified with Phusion Hot Start Flex 2X Master Mix (New England Biolabs, Ipswich, MA, USA) using the following primers: PulA-74F, 5’-GTACTTCAGCTATTATGAGTCTTTGG-3’ and PulA-3651R, 5’-GCGCTTTCCATTCTTCTTAACT-3’. Based on the RL03 *L. crispatus pulA* sequence, this assay generates a 3578 bp product. Select products were sequenced using the Sanger method by the Centre for Health Genomics and Informatics (CHGI) at the University of Calgary and analyzed in SnapGene®. To interrogate the location, type and predicted functional consequences of *pulA* mutations, an alignment of each *pulA* sequence containing a mutation, along with the RL03 reference sequence, was created in SnapGene® using MUSCLE 3.8.1551 with default parameters.

### Phylogenetic analyses

To assess phylogenetic relationships amongst *L. crispatus* genomes and MAGs, *cpn60* phylogenetic marker gene sequences were extracted from Prokka [82] genome annotation files using the gene product identifier “60 kDa chaperonin” and coordinates. Multiple sequence alignment of 115 full-length *L. crispatus cpn60* sequences and a *Lactobacillus acidophilus cpn60* sequence included as the outgroup (strain La-14, NCBI accession GCF_000389675.2:NC_021181.2) was performed using MUSCLE [83] with default parameters. A maximum likelihood (ML) tree was constructed in RAXmL using the GTRCAT model for nucleotide sequence alignments [84, 85]. Branches with less than 50% reproducibility over 1000 bootstrap iterations were collapsed [86].

### Statistical analyses

To account for differences in swab load, all biomolecule measurements were normalized to the total protein content of each swab supernatant or cell pellet (ng). Measurements that fell below the assay limit of detection were replaced with a value equal to half the lower limit of the standard curve. Differences in biomolecule abundance or activity across sample groups were assessed with a Student’s t-test or a Mann-Whitney U-test. Relationships between biomolecules were assessed with a generalized linear regression model and Spearman’s correlation. Statistics were performed in GraphPad Prism (GraphPad, San Diego, CA) or Stata 15 (StataCorp, College Station, TX) and graphs were prepared in GraphPad Prism or R Studio [61]. GraphPad was used to assess data variance (F-test) and normality (Shapiro-Wilk test), and statistical significance was evaluated with an alpha value ≤0.05 and the following significance levels: *p≤0.05, **p≤0.01, ***p≤0.001, ****p≤0.0001.

## Results

### Behavioral, clinical and socio-demographic characteristics of a pilot cohort of Kenyan adolescent girls and young women (AGYW)

Seventeen consenting AGYW who attended the baseline visit of a prospective sexual health study in Thika, Kenya [54], but did not subsequently return to the clinic, were selected for our pilot study exploring new methods of detecting glycogen and related enzyme activities in vaginal swab samples. Clinical data from their baseline assessment, along with behavioral and socio-demographic characteristics reported via questionnaire are summarized in Table 1 (see also Table S1, Additional File 2). The median age of participants in the pilot cohort was 18.8 years (IQR 16.4–20.7) and the median years of schooling was 12 (IQR 10.5–12). Compared to the full cohort of 400 participants [54], fewer participants in the pilot cohort resided in homes with running water (29% pilot vs. 48% full), electricity (35% pilot vs. 69% full) and concrete floors (59% pilot vs. 73% full). Frequency of reported sexual debut (20–24%), median age of sexual debut (18) and median number of sexual encounters in the last three months (1) were similar in the pilot and full study cohorts. All 17 pilot participants (100%) were BV-negative at baseline and 1 participant (6%) tested positive for an STI (trichomoniasis).

### Vaginal microbiome composition in Kenyan AGYW

Taxonomic profiles of the vaginal microbiome in pilot study participants were generated using shotgun metagenomic sequencing. Considering a definition of dominant species as those comprising ≥50% relative abundance [87, 88], each sample was dominated by one or two species. *Lactobacillus* species dominated the vaginal niche in 14 samples (82%), with dominant *L. crispatus* observed in 11 samples (65%; Figure 1, Tables S4–S5, Additional File 2). Two samples (12%) had *L. iners-*dominated vaginal microbiotas and one sample (6%) had a *L. johnsonii-*dominated microbiota. *Gardnerella* species were dominant in two samples (12%), while *Bifidobacterium breve* was dominant in one sample (6%). Sexual debut was reported in 4 participants (24%), all of which exhibited a *Lactobacillus-*dominated microbiome (*L. crispatus* N=3; *L. iners* N=1). Although all pilot participants presented with low Nugent scores (0–4; BV-negative), the highest Nugent score was observed in a *Gardnerella-*dominated sample. One participant had yeast (*Candida*) detected in their metagenome; this participant and another experiencing trichomoniasis both had *L. crispatus-*dominated microbiomes. Days since last menstrual period and age showed no clear relationship with the composition of the vaginal microbiome and were not significantly different between samples dominated by *L. crispatus* versus other taxa (days since last menstrual period: Mann-Whitney U-test ns p=0.713, age: Student’s t-test ns p=0.394).

**Figure 1.**
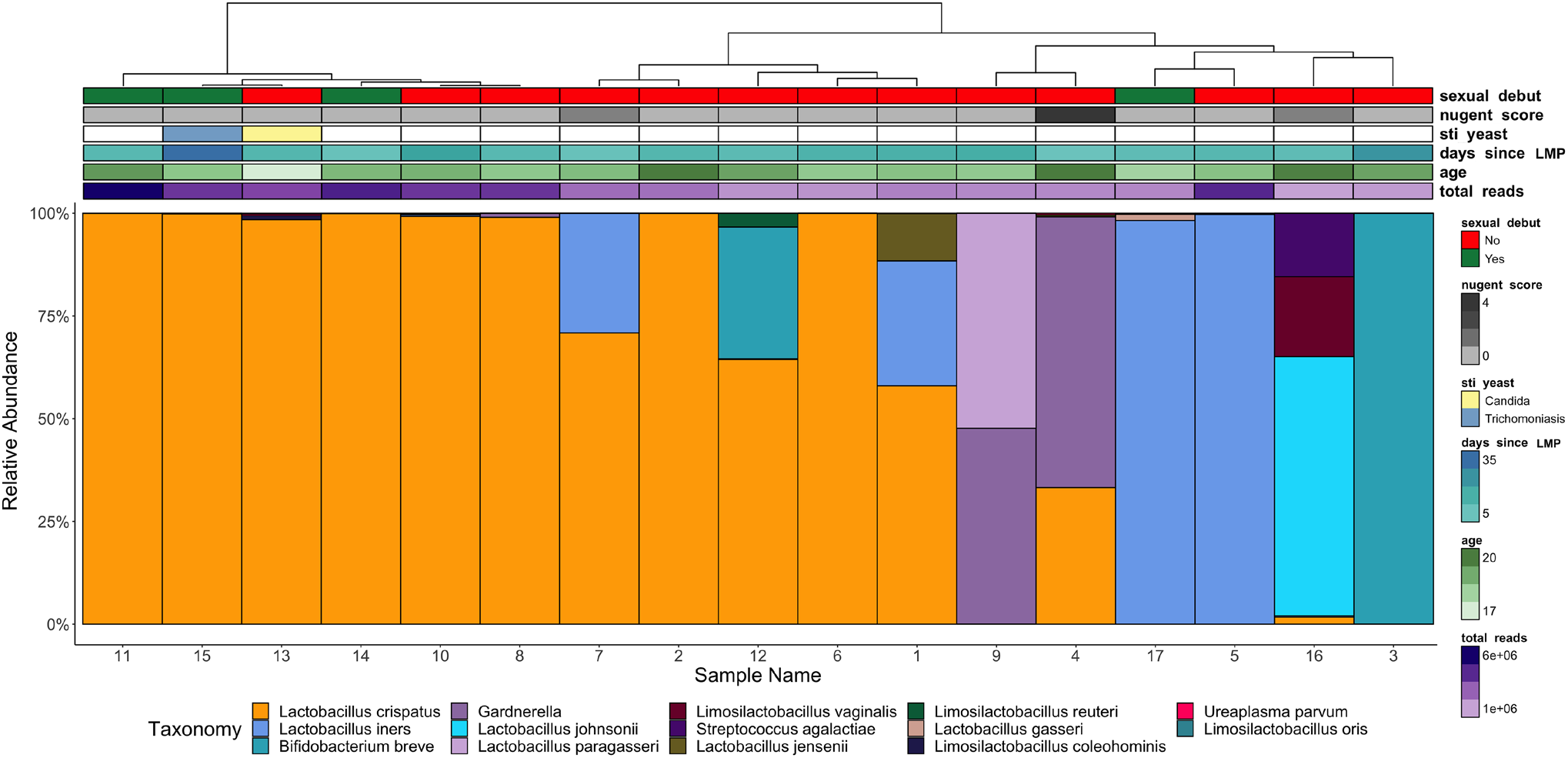
Species-level composition of the vaginal microbiome in Kenyan adolescent girls and young women (AGYW). Hierarchical clustering and bacterial taxonomic relative abundance profiles of the vaginal microbiota from the pilot cohort of Kenyan AGYW (N=17). Only taxa classified as bacteria with relative abundance greater than 0.01% are shown. Metadata categories including sexual debut, Nugent score (BV-negative: 0*–*3, BV-intermediate: 4*–*6, BV-positive: ≥7), detection of sexually transmitted infections (trichomoniasis) or yeast (*Candida*), days since last menstrual period (LMP), age and total metagenomic reads are overlayed above the taxonomic profiles for each participant.

### Differential detection of glycogen and glycogen-derived maltodextrin highlights unique relationships with lactic acid

Previous studies have shown levels of cell-free glycogen in vaginal fluid are positively correlated with *Lactobacillus* spp. load and inversely correlated with pH [41, 88]. This is somewhat counterintuitive, however, since higher loads of glycogen-degrading *L. crispatus* would presumably deplete glycogen as glycogen catabolites (i.e. maltodextrin, maltotriose, maltose) accumulate. We examined whether this incongruity could be explained by nonspecific measurement of glycogen in complex human vaginal samples. Fluorometric methods of glycogen quantification [41, 42, 51, 88–90] apply glucoamylase enzymes to hydrolyze glycogen to glucose, which is then fluorometrically measured. We confirmed that these fluorometric methods also hydrolyze and detect glycogen catabolites, including maltodextrin, maltotriose and maltose (Figure S1, Additional File 1). Thus, to specifically measure glycogen in human vaginal samples, we applied a size exclusion approach that takes advantage of glycogen’s large size, estimated at 10^7^ kDa, and unique highly branched structure [43]. From a mixture of carbohydrates, glycogen and maltodextrin were differentially detected by the fluorometric method in centrifugally separated high molecular weight (>100 kDa) and low molecular weight (<100 kDa) fractions, respectively (Figure S1, Additional File 1). Glucose was also distinguishable from maltodextrin/maltose in the low molecular weight fraction when hydrolysis enzymes were not added. We applied this method to vaginal swab supernatants, correcting the resulting carbohydrate measurements for differences in swab load by normalizing to the total protein content of each supernatant, which we found correlated with other indicators of vaginal swab load/biomass, namely total bacterial abundance (broad-range 16S rRNA gene qPCR applied to DNA extracted from swab cell pellets; ρ=0.628 **p=0.0083; Figure S2A–C, Additional File 1) and the menstrual cycle (days since last menstrual period, a correlate of epithelial exfoliation rate; ρ=-0.752 *p=0.0011; Figure S2D–E, Additional File 1). Our results showed that the majority of glucose released through the hydrolysis of D-glucose polymers in vaginal swab supernatants was in fact derived from glycogen. Glucose derived from glycogen was 3-fold more abundant, on average, than that derived from maltose, maltotriose and longer α-1,4 glucose polymers, which we collectively refer to as maltodextrin (Wilcoxon signed-rank test p=0.007; Figure 2A). However, in four samples (Samples 1, 3, 9, 13; 23.5%), maltodextrin comprised a majority or all of the D-glucose polymers we detected. No significant difference was observed in the abundance of glycogen or maltodextrin in *L. crispatus-*dominated samples compared to samples dominated by other taxa (glycogen ns p=0.961, maltodextrin ns p=0.256; Figure 2B–C). Next, we evaluated whether glycogen and maltodextrin correlated with fermentative metabolic products: D-, L- or total lactic acid. D-lactic acid levels were significantly higher in samples dominated by *L. crispatus* (*p=0.0202; Figure S3A–B, Additional File 1), while total lactic acid and L-lactic acid levels were similar between *L. crispatus-*dominated and non *L. crispatus-*dominated samples (L-lactic acid ns p>0.999, total lactic acid ns p=0.301; Figure S3A,C–D, Additional File 1). Glycogen showed a positive correlation with all forms of lactic acid, though only relationships with L-lactic acid (ρ=0.591 *p=0.014; Figure 2E) and total lactic acid (ρ=0.598 *p=0.013; Figure 2F) achieved statistical significance. In contrast, maltodextrin exhibited non-significant negative relationships with each lactic acid isomer (Figure 2G–I).

**Figure 2.**
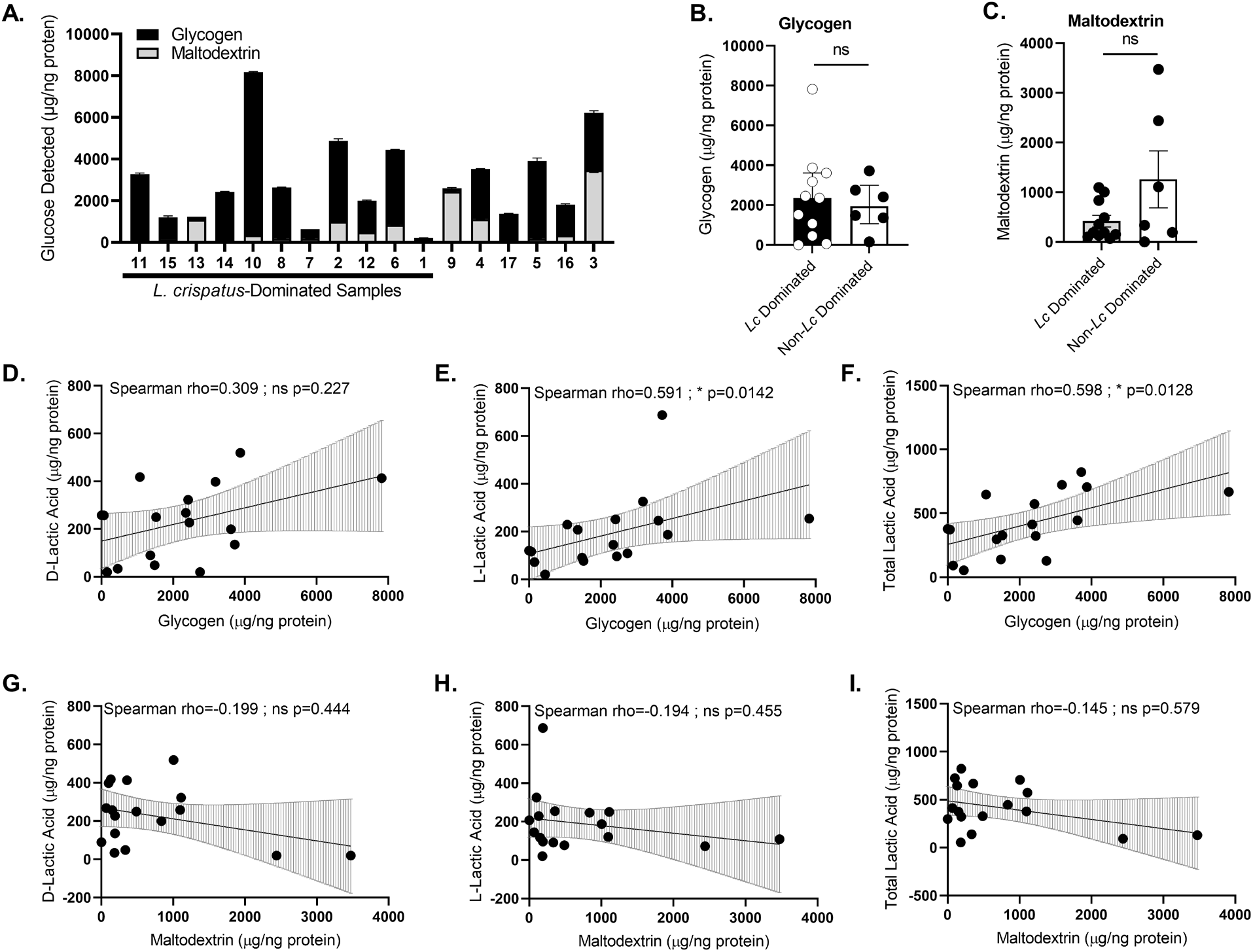
Glycogen is positively correlated with lactic acid levels in vaginal swab supernatants. **(A)** Glycogen and maltodextrin were measured in vaginal swab supernatants collected from Kenyan adolescent girls and young women (N=17). Total glucose derived from glycogen or maltodextrin was normalized to total protein concentration and presented as mean µg carbohydrate/ng of protein ± range from one independent experiment run in technical duplicate. Distribution of **(B)** glycogen and **(C)** maltodextrin in *L. crispatus*-dominated (*Lc* Dominated; N=11) and non-*L. crispatus*-dominated (non-*Lc* Dominated; N=6) vaginal samples presented as median ± interquartile range. Statistical significance of the difference between *Lc* Dominated and non-*Lc* Dominated samples was assessed with a Mann-Whitney U-test for glycogen (ns p=0.961) and maltodextrins (ns p=0.256). **(D–I)** Scatterplots of D-lactic acid, L-lactic acid and total lactic acid measurements as a function of **(D–F)** glycogen or **(G–I)** maltodextrin measurements. Regression lines were fitted using a generalized linear model and error bars indicate 95% confidence intervals. Relationships were assessed with Spearman’s correlation, with the rho value and p-value indicated above each scatterplot.

### Amylopullulanase and amylase activity correlate with glycogen-derived maltodextrin

Bacteria catabolize glycogen via the concerted activity of individual or multifunctional enzymes cleaving linear α-1,4 and branched α-1,6 glycosidic linkages. Human and microbial α-amylases and α-glucosidases cleave α-1,4 linkages, while microbial pullulanase enzymes specifically target α-1,6 linkages [44, 91, 92]. Bacteria can also produce amylopullulanases, multi-functional enzymes capable of cleaving both α-1,4 and α-1,6 glycosidic linkages [93]. We confirmed that a commercially available fluorescent starch degradation assay detects the activities of both pullulanase and amylase enzymes; this assay may therefore be considered a measure of the total ‘amylopullulanase’ glycogen degrading capacity (Figure S4, Additional File 1). When applied to Kenyan AGYW swab samples, this assay detected amylopullulanase activity primarily in swab supernatants, with no significant difference in activity between *L. crispatus*-dominated and non *L. crispatus-*dominated samples (ns p=0.256; Figure 3A,D, Tables S2–S3, Additional File 2). Similarly, α-amylase activity was exclusively detected in swab supernatants and showed no relationship with *L. crispatus* dominance (ns p=0.733; Figure 3B,D, Tables S2–S3, Additional File 2). Intriguingly, amylopullulanase and α-amylase activities did not exhibit any relationship with glycogen (amylopullulanase: ρ=0.0882 ns p=0.734; Figure 3G, α-amylase: ρ=-0.0515 ns p=0.846; Figure 3H), but were strongly correlated with maltodextrin levels (amylopullulanase: ρ=0.829 ****p<0.0001; Figure 3J, α-amylase: ρ=0.664 **p=0.0046; Figure 3K).

**Figure 3.**
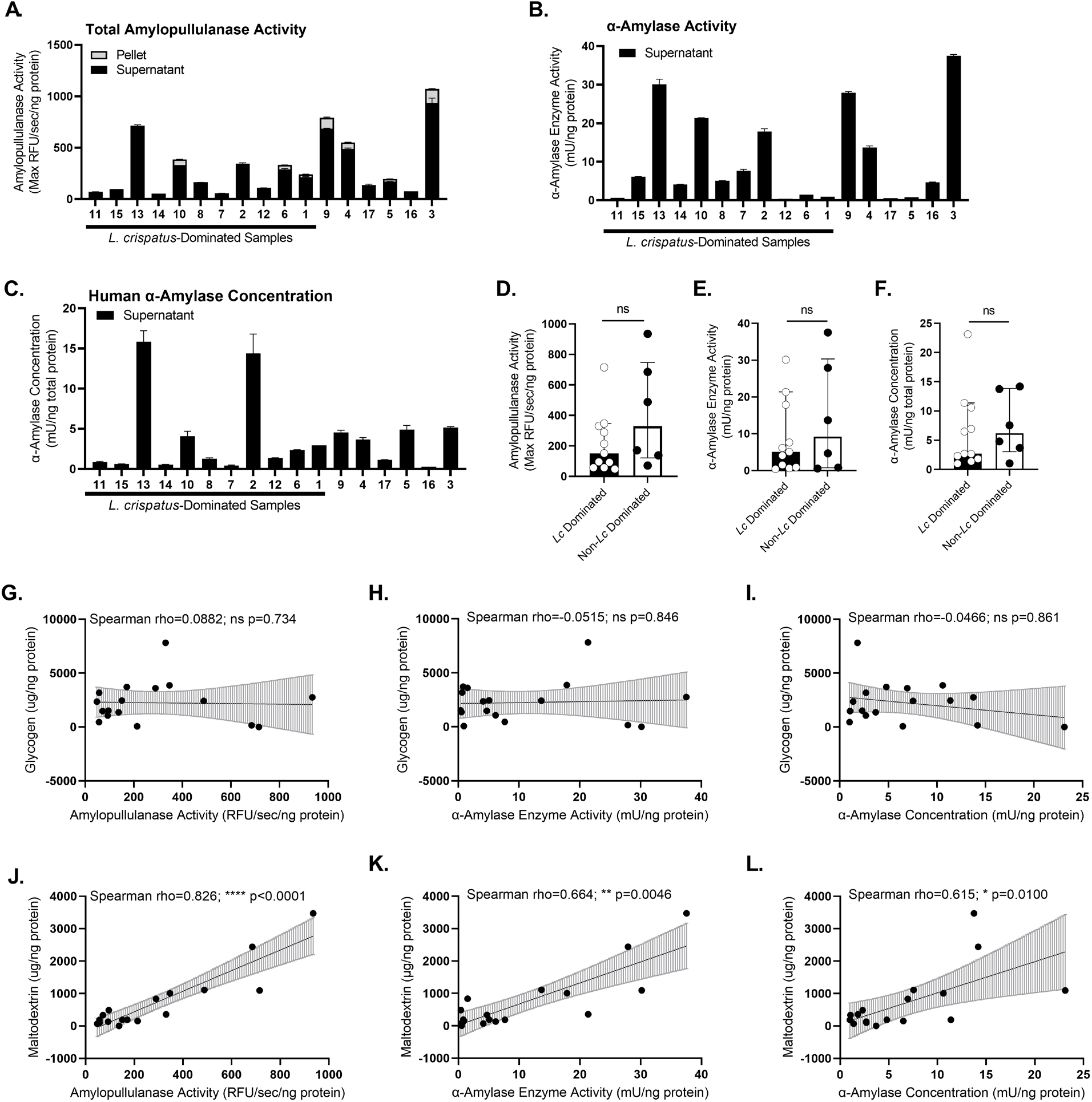
Vaginal amylopullulanase activity is positively correlated with glycogen-derived maltodextrin. **(A)** Total amylopullulanse activity (amylase, pullulanase and amylopullulanase activity), **(B)** α-amylase activity and **(C)** human α-amylase concentration measured in vaginal swab supernatants from Kenyan adolescent girls and young women (N=17). Results were normalized to total protein concentration and presented as mean ± range of **(A)** amylopullulanase activity (max relative fluorescence units (RFU)/seconds/ng protein), α-amylase activity (mU/ng protein) or **(C)** human α-amylase concentrations (mU/ng protein) from one independent experiment run in technical duplicate. **(D–F)** Distribution of **(D)** amylopullulanase activity, **(E)** α-amylase activity and **(F)** human α-amylase concentration in *L. crispatus*-dominated (*Lc* Dominated, N=11) and non-*L. crispatus-*dominated (non-*Lc* Dominated, N=6) vaginal samples presented as median ± interquartile range. Statistical significance between *Lc* Dominated and non-*Lc* Dominated samples was assessed with Mann Whitney U-test for amylopullulanase activity (ns p=0.256), α-amylase activity (ns p=0.733) and human α-amylase concentration (ns p=0.462) **(G–L)** Scatterplots of glycogen and maltodextrin levels as a function of **(G,J)** amylopullulanase activity, **(H,K)** α-amylase activity and **(I,L)** human α-amylase concentration. Regression lines were fitted using a generalized linear model and error bars indicate 95% confidence intervals. Relationships were assessed with Spearman’s correlation, with the rho coefficient and p-value indicated above each scatterplot.

Human α-amylase enzyme concentration was positively correlated with amylopullulanase activity (ρ=0.831 ****p<0.0001; Figure S5A, Additional File 1), and trended toward correlation with α-amylase activity (ρ=0.458 ns p=0.0661; Figure S5B, Additional File 1). Relationships of α-amylase enzyme with glycogen and maltodextrin mirrored those seen with the amylase activity assays (glycogen: ρ=0.047 ns p=0.861; Figure 3I, maltodextrin: ρ=0.615 *p=0.01; Figure 3L), and no difference in α-amylase levels was detected across *L. crispatus*-dominated and non *L. crispatus-*dominated samples (ns p=0.462; Figure 3F). Since the additive amylopullulanase assay did not detect α-glucosidases (Figure S4, Additional File 1), an additional assay was performed to quantify α-glucosidase activity in vaginal swab supernatants and cell pellets (Figure S6A, Additional File 1; Tables S2–S3, Additional File 2). This activity exhibited no linkage to *L. crispatus* dominance or glycogen (ns p=0.884; Figure S6B–C, Additional File 1), but showed a trend of positive, but non-statistically significant correlation with maltodextrin levels (ρ=0.439 ns p=0.08; Figure S6D, Additional File 1). A positive correlation was also observed between human α-amylase concentration and α-glucosidase activity in swab supernatants (ρ=0.598 *p=0.0128; Figure S5C, Additional File 1).

### Pullulanase enzyme activity is elevated in *L. crispatus-*dominated samples and correlates with D-lactic acid in a *pulA*-dependent manner

Bacterial pullulanase activity, which cleaves branched glycogen α-1,6 linkages, was evident in most vaginal swab supernatants collected from Kenyan AGYW (Figure 4A), but was not detectable in any of the vaginal swab cell pellet samples tested (Table S3, Additional File 2). Notably, pullulanase activity was 4-fold higher in *L. crispatus-*dominated samples compared to samples dominated by other taxa (*p=0.0273; Figure 4A–4C). In contrast to the other enzymes tested, pullulanase activity did not have a significant relationship with either glycogen or maltodextrin (glycogen: ρ=0.245 ns p=0.342, maltodextrin: ρ=-0.07 ns p=0.773; Figure S7, Additional File 1). Presence of the *L. crispatus* type I pullulanase (*pulA*) gene in Kenyan AGYW samples was evaluated by performing a *pulA* BLASTn search against each metagenome, and confirmed through metagenomic read coverage analysis and a *pulA* gene-targeted PCR assay. These inquiries revealed that *L. crispatus pulA* was absent in 8/17 samples (47%), including 3/11 *L. crispatus*-dominated samples (27%; Figure 4B, Figures S8–S9, Additional File 1, Tables S6–S9, Additional File 2). Each *L. crispatus pulA* gene detected in Kenyan AGYW metagenomes was free of insertions/deletions or nonsense mutations, with the exception of that identified in *L. crispatus*-dominated sample 14, which was found to possess a single nucleotide polymorphism (G514T) that introduces a premature stop codon at amino acid 172 (Table 2, Figure 5A, Figure S9B, Additional File 1). The fact that this mutation is predicted to severely truncate the normally 1259 amino acid PulA protein corresponds with sample 14 possessing less than half of the pullulanase activity detected in *L. crispatus*-dominated samples with an intact *pulA* gene (Figure 4A– 4B). Altogether, pullulanase activity was 66% lower in *L. crispatus*-dominated samples with an absent or inactivated *pulA* gene (4/11 samples, 36%) compared to *L. crispatus-*dominated samples predicted to encode a functional gene (**p=0.0061; Figure 4D). Median *L. crispatus* abundance (read count) was also approximately five-fold lower in *L. crispatus*-dominated samples when *pulA* was absent (active vs. absent/inactive *pulA*: ns p=0.131, present vs. absent *pulA*: *p=0.0121; Figure 4E). Glycogen and maltodextrin levels in *L. crispatus*-dominated samples were not impacted by the presence of a functional *pulA* gene (Figure S10, Additional File 1), but there were downward trends in median D-lactic acid (36% ns p=0.257) and total lactic acid (39% ns p=0.450) levels in *L. crispatus*-dominated samples lacking a functional *pulA* gene (Figure 4F–H). Complementing these observations, pullulanase activity exhibited a significant positive relationship with D-lactic acid and total lactic acid, but not L-lactic acid (D-lactic acid: ρ=0.630 **p=0.008, total lactic acid: ρ=0.532 *p=0.030; Figure 4I– K). This relationship with lactic acid appears to be specific to pullulanase activity, as amylopullulanase activity, α-amylase activity, human α-amylase enzyme concentration and α-glucosidase activity were not correlated with D-, L- or total lactic acid (Figure S11, Additional File 1). Further demonstrating the distinct contribution of pullulanases, their activity was not significantly correlated with amylopullulanase or α-amylase activity, although these activities were significantly correlated to each other (Figure S12).

**Figure 4.**
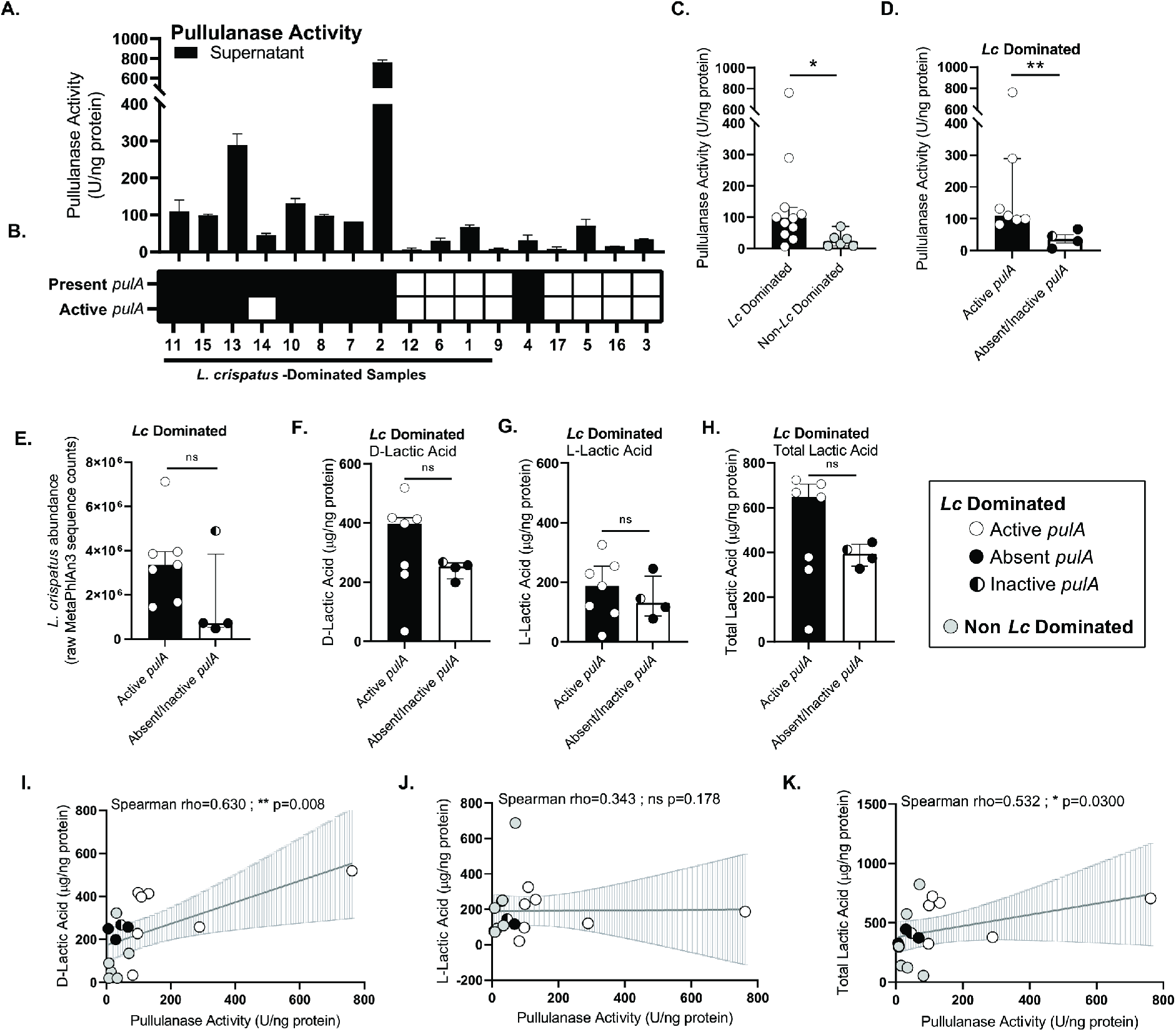
*L. crispatus*-dominated vaginal samples have elevated pullulanase activity that correlates with D-lactic acid levels. **(A)** Pullulanase enzyme activity measured in vaginal swab supernatants collected from Kenyan AGYW (N=17). Results were normalized to total protein concentration and presented as mean U/ng protein ± range with the y-axis split at 400 from one independent experiment run in technical duplicate. **(B)** Presence of the *L. crispatus pulA* gene in Kenyan AGYW samples. *pulA* gene presence and predicted activity was assessed in metagenomes and through targeted PCR as described in the Methods and Table 2 footnotes. Present/active *pulA* genes are indicated in black, while absent/inactive *pulA* genes are indicated in white. **(C)** Distribution of pullulanase activity in *L. crispatus*-dominated (*Lc* Dominated; N=11) and non-*L. crispatus*-dominated (non-*Lc* Dominated; N=6) vaginal samples presented as median ± interquartile range with the y-axis split at 400. Statistical significance was assessed with a Mann-Whitney U-test (*p=0.0273). **(D)** Distribution of pullulanase activity in *L. crispatus*-dominated samples with predicted active *pulA* genes (N=7) and absent/inactive *pulA* genes (N=4) presented and statistically assessed as described for panel **(C)** (**p=0.0061). **(E)** Distribution of *L. crispatus* raw MetaPhlAn3 sequence counts in samples dominated by *L. crispatus* with predicted active *pulA* gene (N=7) and absent/inactive *pulA* gene (N=4) presented as median ± interquartile range. Mann-Whitney U-tests were used to assess statistical significance (present/active *pulA* vs. absent/inactive *pulA*: ns p=0.164, present (regardless of activity) *pulA* vs. absent *pulA* gene: *p=0.0121). **(F–H)** Distribution of **(F)** D-lactic acid, **(G)** L-lactic acid and **(H)** total lactic acid levels (µg/ng protein) in *L. crispatus*-dominated samples with predicted active *pulA* gene (N=7) and absent/inactive *pulA* gene (N=4) presented as median ± interquartile range. Mann-Whitney U-tests were used to assess statistical significance across indicated *pulA* groups (D-lactic acid: ns p=0.315, L-lactic acid: ns p=0.649, total lactic acid: ns p=0.527). **(I–K)** Scatterplots of **(I)** D-lactic acid, **(J**) L-lactic acid and **(K)** total lactic acid as a function of pullulanase activity. Regression lines were fitted using a generalized linear model with error bars indicating 95% confidence intervals. Relationships were assessed using Spearman’s correlation, with the rho coefficient and p-value indicated above each scatterplot.

**Figure 5.**
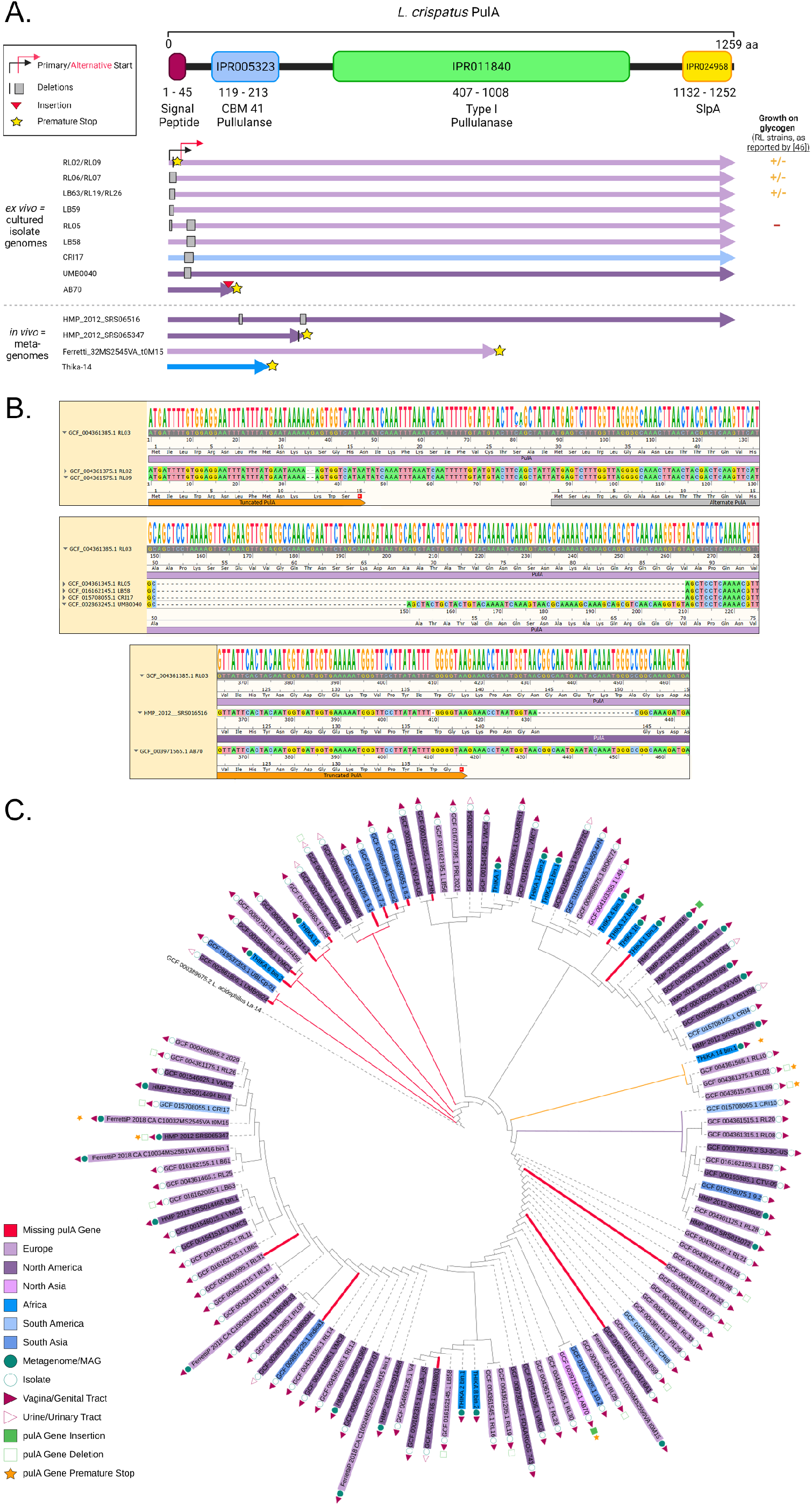
Diverse *L. crispatus pulA* mutations show wide geographic distribution, while gene loss is enriched in select clades. **(A)** Schematic illustration of *L. crispatus* PulA protein domain architecture juxtaposed against the location and type of genetic mutations observed in *L. crispatus pulA* alleles assembled in our analysis of 86 bacterial isolate genomes (*ex vivo*) and 29 vaginal metagenomes (*in vivo*). The size and location of deletions, the most common type of mutation, are represented by black lines (1–3 nucleotides) and grey boxes (>3 nucleotides, approximate to size). Location of the alternate start predicted by van der Veer et al. [46] to result in intracellular localization of functional PulA protein in strains with N-terminal deletions is shown with a jointed red arrow. Deletions, insertions or single nucleotide polymorphisms (SNPs) resulting in a premature stop codon are indicated with a yellow star at the predicted point of truncation. The color of each *pulA* allele represents the geographic origin of that strain/sample, as shown in **(C)**. Figure was created with Biorender.com. **(B)** SnapGene® visualizations of a *pulA* MUSCLE alignment highlighting the linkage between diverse genetic mutations and predicted changes to the protein sequence. Each box highlights a different segment of the full-length *pulA* gene alignment with the nucleotide consensus (>50%) at the top, followed by the RL03 reference gene sequence (grey) and select mutant gene sequences (nucleic acid color coding). **(C)** Phylogenetic tree of *cpn60* gene sequences obtained from 115 *L. crispatus* isolates and vaginal metagenomes that produced *L. crispatus* metagenome assembled genomes (MAGs). Strains/metagenomes lacking a *pulA* gene sequence are highlighted as thick red leaves. Clades with ≥50% of strains/metagenomes lacking a *pulA* gene sequence are indicated by thin red branches. Clades with ≥50% of strains/metagenomes containing a premature stop codon are indicated by thin yellow branches. Clades containing at least 5 sequences showing no mutations are indicated by think purple branches. All other leaves (dotted lines) and branches (solid lines) are coloured grey.

**Table 2.** Detection and mutation frequency of *Lactobacillus crispatus* type I pullulanase (*pulA*) genes in isolate genomes and vaginal metagenomes

|  | *In vivo*<sup>a</sup> |  |  | *Ex vivo*<sup>b</sup> | **Total *in vivo***<br>**and *ex vivo***<br>N=115<br>n (%) |
| --- | --- | --- | --- | --- | --- |
|  | Kenyan<br>AGYW Pilot<br>N=12<sup>c</sup><br>n (%) | Pasolli et<br>al. [75]<br>N=17<br>n (%) | **Total**<br>N=29<br>n (%) | NCBI<br>Genomes<br>N=86<br>n (%) |  |
| **Present/Full-length Gene** | 8 (66.7) | 15 (88.2) | **23 (79.3)** | **58 (67.4)** | **81 (70.4)** |
| **Absent/Mutant Gene** | 4 (33.3) | 2 (11.8) | **6 (20.7)** | **28 (32.6)** | **34 (29.6)** |
| Absent<sup>d</sup> | 3 (25.0) | 0 (0) | 3 (10.3) | 14 (16.3) | 17 (14.8) |
| N-terminal Deletion<sup>e</sup> | 0 (0) | 0 (0) | 0 (0) | 7 (9.3) | 7 (6.1) |
| Internal Insertion/Deletion<sup>f</sup> | 0 (0) | 1 (5.9) | 1 (3.4) | 7 (9.3) | 8 (7.0) |
| Nonsense Mutation<sup>g</sup> | 1 (8.3) | 1 (5.9) | 2 (6.9) | 0 (0) | 2 (1.7) |

<sup>a</sup> Vaginal metagenomes that produced *L. crispatus* metagenome assembled genomes (MAGs)
<sup>b</sup> Cultured *L. crispatus* isolate genomes obtained from NCBI
<sup>c</sup> *pulA* gene presence/absence was confirmed by gene-targeted PCR using DNA extracted from vaginal swab pellets
<sup>d</sup> Missing genes were defined as having no BLASTn hit returned using the *L. crispatus* RL03 *pulA* query sequence
<sup>e</sup> Genes were considered to have N-terminal deletions if the signal peptide was truncated or missing (N-terminal deletion), potentially disrupting protein secretion
<sup>f</sup> Internal insertions could be in frame or result in a frameshift mutation.
<sup>g</sup> Single nucleotide polymorphisms (SNPs) resulting in a premature stop codon are reported as nonsense mutations. Other SNPs resulting in amino acid substitutions are not reported.

Given the surprising observation that *L. crispatus pulA* genes were commonly absent or inactivated in our Kenyan samples (4/12 metagenomes producing *L. crispatus* MAGs, 33%; Table 2), we sought to examine the frequency of *pulA* gene loss and mutation across a broader range of human samples (“*in vivo*”) and *L. crispatus* isolate genomes (“*ex vivo”*). Using the same BLASTn approach, we found that *pulA* was absent or mutated in 2/17 metagenomes from European and North American women that produced *L. crispatus* MAGs (12%) [75] and 28/86 publicly available *L. crispatus* genomes from cultured genitourinary isolates (33%; Table 2, Tables S10–S12, Additional File 2). We identified a variety of *pulA* mutations, most of which occurred in the first third of the coding sequence (Figure 5A; Table S13, Additional File 2). One of the most common types of mutations were short N-terminal deletions previously reported by van der Veer et al. [46]; these all occurred before a putative alternate start codon (M30; Figure 5B). One or more internal deletions before the type I pullulanase enzymatic domain constituted another common mutation type, with three of these resulting in frameshift mutations that introduced premature stop codons. Nucleotide insertions and single nucleotide polymorphisms (SNPs) contributing premature stop codons were less frequent, with the latter only observed *in vivo* (Figure 5A–B; Table S13, Additional File 2).

Loss of the *pulA* gene was common in *L. crispatus* strains colonizing Kenyan AGYW (3/12 metagenomes producing *L. crispatus* MAGs, 25%), but was not observed in any metagenomes assembled from European or North American women (0/17 metagenomes producing *L. crispatus* MAGs; Table 2; Table S13, Additional File 2). This led us to ask whether *pulA* gene loss was linked to *L. crispatus* strain lineage or geography. Overlaying *pulA* gene presence/absence and mutation type on a *cpn60* gene-based phylogeny of all 115 *L. crispatus* strains (*in vivo* and *ex vivo*) revealed that *pulA* gene loss and mutation were not restricted to a particular lineage (Figure 5C). However, some lineages were enriched for *pulA* gene loss or premature stop codons and only one deeply branched lineage contained no strains with *pulA* gene loss or mutation. Although limited by a small number of strains from most regions, geographical inquiry revealed that *pulA* gene loss was also relatively common in strains from South Asia (5/9 strains, 56%), but more rare in strains from Europe (3/46 strains, 7%) and North America (6/42 strains, 14%).

## Discussion

This pilot study is the first of its kind to link vaginal microbiome features with biomarkers of glycogen utilization and vaginal health in young African women around the time of sexual debut. It was common for the adolescent girls and young women (AGYW) in this small subset of our larger Kenyan cohort to have a vaginal microbiome dominated by *L. crispatus*. As such, the prevalence of *L. crispatus* colonization in this suburban pilot study (76%) was very similar to that in a cohort of 386 17– 18 year-old school girls in urban Tanzania (69%), 42% of which reported sexual debut [28]. In contrast, a cohort from a rural area of Kenya, where socioeconomic indicators like electricity and running water were more rare, exhibited a lower rate of *L. crispatus* dominance, which coincided with a prevalence of BV that was double that in our baseline study population [29]. These well-powered cohorts recruited in Tanzania and rural Kenya both showed a negative impact of sexual debut on *L. crispatus* bacterial load, including *L. crispatus* dominance [29]. Reporting very limited sexual experience, the small number of sexually active participants in our pilot cohort each had a vaginal microbiome dominated by *Lactobacillus* species. Participants lacking LD, with vaginal microbiotas dominated by *G. vaginalis* or *B. breve*, reported no prior sex. Vaginal colonization by BV-associated bacteria has also been seen in AGYW who reported no penile-vaginal sex in other studies [27, 100], and is largely attributed to the well-documented underreporting of sexual activity by AGYW [101]. Ultimately, profiling the vaginal microbiome of our full longitudinal cohort (N=400) [54] will be necessary to understand how sexual debut and increased sexual encounters impact *Lactobacillus* dominance and the susceptibility of Kenyan AGYW to BV.

The high prevalence of *L. crispatus* in this pilot cohort provided a unique opportunity to examine biomolecules associated this keystone species and improve understanding of its ecological stability. Several previous studies observed a positive correlation between ‘cell-free’ glycogen and the abundance of *Lactobacillus* spp. in vaginal fluid, showing varying ability to achieve statistical significance with *L. crispatus* [41, 88], *L. jensenii* [41, 42, 88] and *L. iners* [42]. Since observational studies have suggested that estrogen-dependent glycogen deposition precedes the colonization of pubescent vaginal tissue by lactobacilli [94, 95], one common interpretation of the glycogen-*Lactobacillus* correlation is that glycogen in some way drives, or must be maintained above a critical threshold for LD to occur [41, 42, 44]. Our study, using methods that distinguish glycogen from maltodextrin, found no relationship between either biomolecule and *L. crispatus* dominance, although there appeared to be a trend for maltodextrin levels to be lower in samples with *L. crispatus* dominance. With the limitation of our small sample size, these findings require confirmation in future work.

Our study further revealed that glycogen and maltodextrin exhibit distinct relationships with lactic acid, the former positively correlated and the latter inversely correlated. This linkage of higher glycogen levels with high lactic acid (low pH) supports a hypothesis put forward by Spear et al. and Nunn et al. [88, 96], which posits that the positive correlation between vaginal glycogen levels and *Lactobacillus* abundance is driven by the glycogen degradation activities of bacteria other than lactobacilli when the pH is high. According to this model (Figure 6), dominant *L. crispatus* (and perhaps other lactobacilli) directly access glycogen, but do not deplete it. Theoretically, *L. crispatus* could selfishly utilize glycogen by directly importing the maltodextrin products of its cell surface pullulanase and releasing few ‘free’ maltodextrin catabolites into the milieu. In contrast, when environmental disruption allows the pH to become more permissive (≥5.5), *G. vaginalis*, and perhaps other pullulanase-expressing BV-associated bacteria such as *P. bivia* and *M. mulieris*, efficiently deplete glycogen and release abundant maltodextrins [51, 53, 88, 97]. This supplies the broader community with a more accessible carbohydrate nutrient pool and forces *L. crispatus* into a more competitive mode of acquiring the fermentative substrates it needs to produce lactic acid and regain the upper hand.

**Figure 6.**
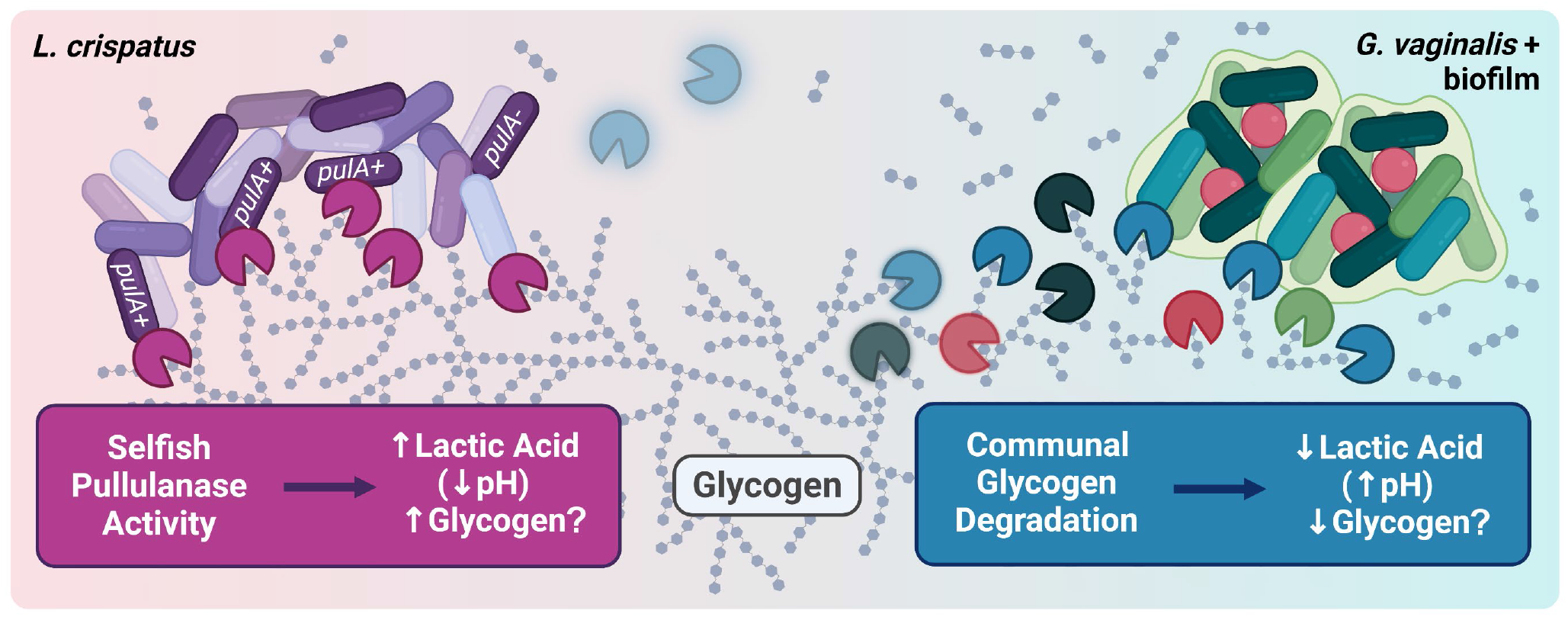
Proposed mechanistic model of the connections between glycogen, glycogen degrading enzymes and lactic acid production in the human vagina. We hypothesize that *L. crispatus* (purple bacteria) may display its pullulanase enzymes (pink Pacmans) as part of a larger glycogen harvesting complex on the cell surface, allowing it to selfishly access glycogen. This is supported by our observation that pullulanase activity does not correlate with increased maltodextrin (linear grey glucose polymers detached from branched glycogen), whereas amylase activity does. Our findings furthermore suggest *L. crispatus* pullulanase activity is important for optimal lactic acid production. By establishing a low pH (pink shading), this activity helps inhibit the growth of *G. vaginalis* and other anaerobic bacteria associated with vaginal biofilms and bacterial vaginosis. Low pH is also expected to inhibit glycogen degrading enzymes (fuzzy blue Pacmans, top center) secreted by the host and bacteria other than lactobacilli; this inhibition potentially underlies the observation that higher glycogen levels can accompany *Lactobacillus* dominance. If the gene encoding pullulanase is lost or mutated (*pulA-*), *L. crispatus*’ nutrient sources would be expected to become more restricted, potentially resulting in a heavy reliance on the maltodextrin released by host amylase and other bacterial glycogen degrading enzymes (red/green/blue Pacmans). This may slow the species’ growth and lactic acid production, especially when more competitive or stressful conditions arise, providing a window for higher pH (blue shading) to activate communal glycogen degradation and the growth of other bacteria (red/green/blue bacteria) that together destabilize *Lactobacillus* dominance.

In line with the above-described model, our data suggest that one of the keys to the ecological success of *L. crispatus* is its pullulanase activity. Upon performing the first quantitative segregation of distinct glycogen-degrading enzyme activities in human samples, we uncovered surprising evidence that pullulanase activity correlates with higher lactic acid production, while amylase activity/concentration and α-glucosidase activity do not. With the commonly-employed fluorometric starch degradation assay, which we describe as detecting amylopullulanase activity (encompassing both pullulanase and amylase activities), the unique relationship between pullulanase activity and lactic acid production was masked. Recent *in vitro* cultivation studies have shown that full length type I pullulanase (*pulA*) alleles with an intact signal peptide sequence are necessary for *L. crispatus* to exhibit robust growth when glycogen is supplied as a sole carbon source [46]; however, *pulA* was not needed for *L. crispatus* growth on glucose and the species has been documented growing on maltose and maltodextrin substrates *in vitro* [98]. Thus, although *L. crispatus* pullulanases may be found at low concentration or exhibit lower activity than amylases *in vivo*, they appear to contribute to the production of D-lactic acid in unique, but as yet poorly understood ways.

Previous work by van der Veer et al. established that *pulA* was variably present and frequently mutated in 28 *L. crispatus* genomes collected from European women [46]. Extending this work to a large set of globally sourced genitourinary *L. crispatus* genomes and vaginal metagenomes revealed that rates of *pulA* gene loss or inactivation are nearly as high *in vivo* as *ex vivo*. Although these findings suggests that *pulA* is not required for *L. crispatus* to dominate the vaginal microbial community in at a cross-sectional moment of time, longitudinal studies are needed to establish whether *pulA(-) L. crispatus* strains display poor stability or resilience over time. Our pilot cohort of Kenyan AGYW also unexpectedly exhibited a higher rate of *in vivo pulA* gene loss/inactivation compared to metagenomes from other parts of the world. However, more extensive phylogenomic analyses are needed to determine whether *pulA* allelic variation or other aspects of *L. crispatus* strain heterogeneity contribute to population-level differences in the prevalence of *L. crispatus* dominance and BV. Though preliminary and based on a very small number of samples, our findings suggest *pulA(-) L. crispatus* strains may exhibit signs of poorer ecological fitness, including lower abundance and reduced lactic acid production, compared to their *pulA(+)* counterparts. A limitation of our study design is that we did not directly measure vaginal pH and thus cannot determine whether the observed changes in lactic acid concentration correlate with transitions across clinically or biologically relevant thresholds of pH. Furthermore, an open question that requires examination with more sensitive and quantitative methods is whether *pulA* tends to be absent in *L. crispatus* strains found at low abundance *in vivo*.

As we pioneered new avenues of biomolecularly profiling the vaginal niche, we encountered several methodological issues that could bias assessment of human samples and contribute to erroneous or inconsistent results. First, like previous studies that normalized vaginal enzyme concentrations and activities to total protein [41, 42, 51, 88, 90], we chose to normalize all biomolecules and activities to total protein to account for the well-documented variability in swab load [31, 51, 99]. As more studies seek to integrate ‘omics, targeted biomolecule assays and clinical metadata, each bringing a unique data structure, limitations/biases and discipline-standard statistical approaches, it will be increasingly important to define ‘best practices’ for data normalization, reporting and statistical inference. Second, the reported confounding of glycogen measurements with maltodextrin illustrates that ‘off the shelf’ biochemical test kits, though accessible and convenient, are not necessarily designed or validated for assaying highly complex biological samples. Well-designed control experiments are needed to ensure methods are appropriately selected and interpreted. Third, having demonstrated significant levels of glycogen-degrading enzyme activity in biobanked vaginal samples, our work underlines the critical importance of cold chain management to ensure measured glycogen/maltodextrin levels reflect those present at the time of sample collection. Finally, in contrast with previous work, we found glycogen-degrading enzyme activities, including that of pullulanases, were abundant in vaginal swab supernatants and minimally present in the cell pellet [97]. Since N-terminal motifs in the *L. crispatus* PulA sequence are predicted to tether the protein to the outer membrane’s Surface Layer Protein A [97], it is possible that most PulA protein is cell surface-associated *in vivo*, but was perhaps dislodged through the vigorous vortexing performed during swab preparation. Alternatively, PulA may be released from the cell wall *in vivo* through biological processes such as proteolytic activity. These uncertainties illustrate the importance of rigorously documenting sample handling methods, and provide impetus for future studies that identify the mechanisms and conditions shaping PulA’s association with the cell wall.

## Conclusions

This exploratory study provided insights into the vaginal microbiome of young African women around the time of their sexual debut, confirming that communities dominated by *L. crispatus* are most common. In seeking to understand the biomolecular drivers of this natural state, we show that although amylase activity is the primary driver of glycogen catabolite accumulation in vaginal fluid, pullulanase activity, contributed predominantly by *L. crispatus* in our samples, appears to maximize D-lactic acid production. Scaling this analysis to our longitudinally collected samples will address whether genomic and enzymatic indicators of *L. crispatus* pullulanase activity are predictive of sustained *Lactobacillus* dominance and vaginal health.

## Additional Files

**Additional File 1: Supporting figures (DOC). Figure S1**. Validation of a new method for differential detection of glycogen, maltodextrin and glucose in vaginal samples. **Figure S2**. Correlates of total protein in vaginal swab supernatants. **Figure S3**. Elevated D-lactic acid levels in vaginal samples dominated by *L. crispatus.* **Figure S4**. Enzyme activities detected in a fluorescent starch degradation assay. **Figure S5**. Human α-amylase correlates with amylopullulanase and α-glucosidase activities in vaginal swab supernatants, but not pullulanase activity. **Figure S6**. α-glucosidase activity correlates with maltodextrin levels in vaginal swab supernatants. **Figure S7**. Pullulanase activity does not correlate with glycogen or maltodextrin levels in vaginal swab supernatants. **Figure S8**. Coverage of the *L. crispatus pulA* gene and flanking regions in vaginal metagenomes dominated by *L. crispatus.* **Figure S9**. *L. crispatus pulA* gene amplification from vaginal swab samples. **Figure S10**. Glycogen and maltodextrin levels do not depend on whether *L. crispatus* encodes a functional pullulanase. **Figure S11**. Amylase and α-glucosidase activities do not correlate with vaginal lactic acid. **Figure S12**. Correlations of enzyme activity in vaginal swab supernatants.

**Additional File 2: Cohort and analysis data used in this study (XLSX)**.**Table S1**. Participant clinical and demographic metadata. **Table S2**. Biomolecule measurements in vaginal swab supernatants. **Table S3**. Biomolecule and bacterial abundance measurements in vaginal pellets. **Table S4**. Raw MetaPhlAn3 bacterial species read counts. **Table S5**. Barcode hopping-adjusted MetaPhlAn3 bacterial species read counts. **Table S6**. Thika metagenome BLASTn results. **Table S7**. Thika MAG BLASTn results. **Table S8**. *pulA* gene coverage in Thika metagenomes. **Table S9**. *Lactobacillus crispatus* genome coverage in Thika metagenomes. **Table S10**. Pasolli metagenome BLASTn results. **Table S11**. Pasolli MAG BLASTn results. **Table S12**. NCBI *L. crispatus* whole genome sequence BLASTn results. **Table S13**. *Lactobacillus crispatus pulA* allele data.

**Additional File 3: Reference information for analysis of genome, metagenome and metagenome assembled genome sequences (XLSX)**. **Table S14**. Thika metagenomic assemblies. **Table S15**. Thika MAGs. **Table S16**. Pasolli metagenomes. **Table S17**. Pasolli MAGs. **Table S18**. NCBI *Lactobacillus crispatus* genomes. **Table S19**. Positional *Lactobacillus crispatus pulA* gene coverage in Thika metagenomes with *a pulA* BLASTn hit. **Table S20**. Positional *Lactobacillus crispatus* genome coverage in Thika metagenomes without a *pulA* BLASTn hit. **Table S21**. Code and algorithm parameters.

## List of Abbreviations

AGYW: adolescent girls and young women
BLASTn: basic local alignment search tool (nucleotide)
bp: base pairs
BV: bacterial vaginosis
GTDB-tk: genome taxonomy database toolkit
HIV: human immunodeficiency virus
HSV: herpes simplex virus
ID: identifier
LD: *Lactobacillus* dominance
LMP: Last menstrual period
MAG: metagenome-assembled genome
Min: minutes
MWCO: molecular weight cut-off
NCBI: National Center for Biotechnology Information
OD: optical density
PCR: polymerase chain reaction
qPCR: quantitative PCR
RA: relative abundance
RFU: relative fluorescence units
SNP: single nucleotide polymorphism
STI: sexually transmitted infection

## Declarations

### Ethics approval and consent to participate

Study approval was obtained from the Kenya Medical Research Institute Scientific Ethics Review Unit (Protocol No. 2760), the University of Washington Institutional Review Board (Study ID STUDY00000946) and the Conjoint Health Research Ethics Board (CHREB) at the University of Calgary (Ethics ID REB18-1832). Each participant provided written informed consent with their guardian present before participating in the study.

### Availability of data and material

All data generated and/or analyzed for the current study are included in the published article and supplementary information files. Patient clinical and demographic metadata, biomolecule measurements and bacterial relative abundance can be found within Additional File 2, Tables S1–S5. Data from *pulA* and metagenomics inquiries are provided in Additional File 2, Tables S6–S13. Reference information for all genome, metagenome and metagenome assembled genomes (MAGs) used in this study can be found in Additional File 3, Tables S14–S20. Algorithm parameters can be found within Additional File 3, Table S21. Cultured isolate genomes were downloaded from the NCBI unfiltered prokaryotes list at: https://www.ncbi.nlm.nih.gov/genome/browse/#!/prokaryotes/ (Accessed August, 2021). Custom snakemake pipelines for adapter trimming, quality filtering, and human sequence decontamination of metagenomic reads, and for running MetaPhlAn3 are available in GitHub: https://github.com/SycuroLab/metqc, https://github.com/SycuroLab/metannotate. R analysis code for the generation of the pheatmap is available here: https://github.com/SycuroLab/Thika-Pilot-AGYW-Repository.

### Authors’ contributions

L.K.S. conceived the hypothesis and experimental design in collaboration with A.C.R. and K.V.L. A.C.R. and N.L.M. designed and oversaw the clinical study and L.O. collected the human samples. K.V.L. and A.C. carried out carbohydrate and enzyme activity assays. S.K. conducted molecular assays including DNA extraction, shotgun library preparation and PCR. K.M. performed metagenomics analysis and *pulA* allele studies. K.V.L., A.C., K.M., S.K. and L.K.S. performed statistical analyses, prepared figures and wrote the manuscript. All authors read, revised and approved of the final version of the manuscript.

## Acknowledgements

The authors would like to thank the adolescent girls and young women who participated in the study, Drs. Wade Abbot, Marc Strous, Lara Mahal, and Lisa Willis for their advice and feedback on the study’s design and interpretation, as well as Megan Kinzel and Stephanie Besoiu for their assistance in our bioinformatics inquiries.

## Additional File 1: Supporting Figures

**Figure S1.**
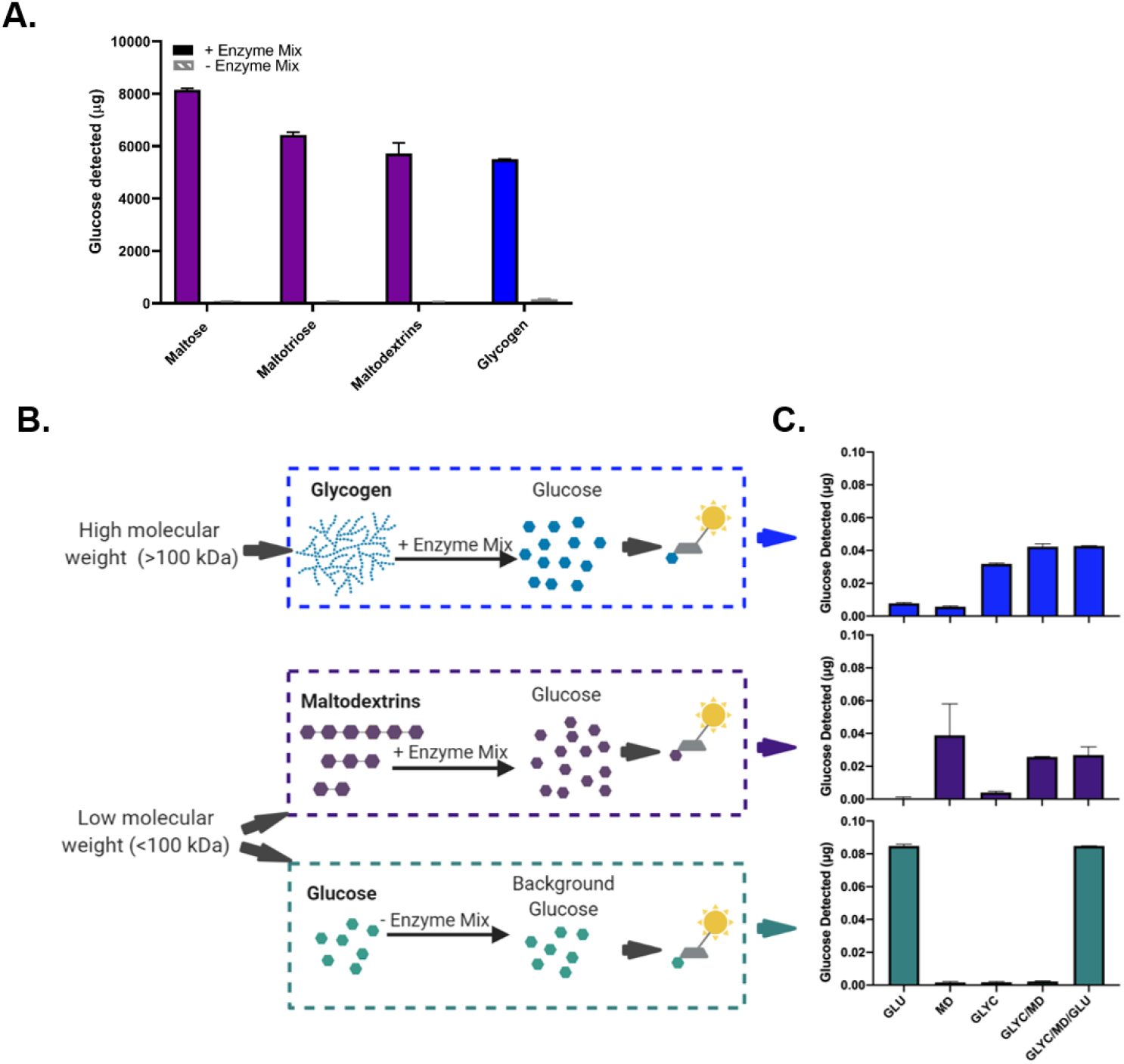
Validation of a new method for differential detection of glycogen, maltodextrin and glucose in vaginal samples. **(A)** Fluorometric glycogen quantification assay (BioVision) detects glucose derived from 3.2 µg/mL solutions of maltose, maltotriose, maltodextrin and glycogen in the presence of the hydrolysis enzyme mix that catalyzes the production of glucose from alpha-glucan (glucose) polymers (solid bars). Control samples without hydrolysis enzyme mix (hatched bars) indicate the background glucose levels, which are subtracted from hydrolyzed samples to determine the concentration of glucose polymers. Results are presented as mean ± standard deviation of 3 technical replicates**. (B)** Schematic illustration of the differential glycogen and maltodextrin quantification assay. Glycogen (>100 kDa) is separated from maltodextrin and glucose (<100 kDa) by size filtration using a 100 kDa molecular weight cutoff centrifugal filter unit. The concentrate fraction is subjected to enzyme hydrolysis to fluorometrically detect glucose derived from glycogen and the filtrate fraction is identically hydrolyzed to derive glucose from maltodextrin (along with maltose and maltotriose). Control samples of the concentrate and filtrate without hydrolysis enzyme mix account for the background glucose levels in each fraction. **(C)** Quantification of glucose (GLU) or glucose derived from solutions of maltodextrin (MD), glycogen (GLYC), a mixture of glycogen and maltodextrins (GLYC/MD), or a mixture of glycogen, maltodextrins and glucose (GLYC/MD/GLU), using solutions containing 3.2 µg/mL of each carbohydrate. Glucose enzymatically released in the high molecular weight concentrates is shown in the top graph (blue, glycogen), while that released in the low molecular weight filtrates is shown in the middle graph (purple, maltodextrin); glucose detected in the absence of the hydrolysis enzyme mix is shown in the bottom graph (teal, glucose). Results are presented as mean ± standard deviation of 3 technical replicates. Schematic illustration created with Biorender.com.

**Figure S2.**
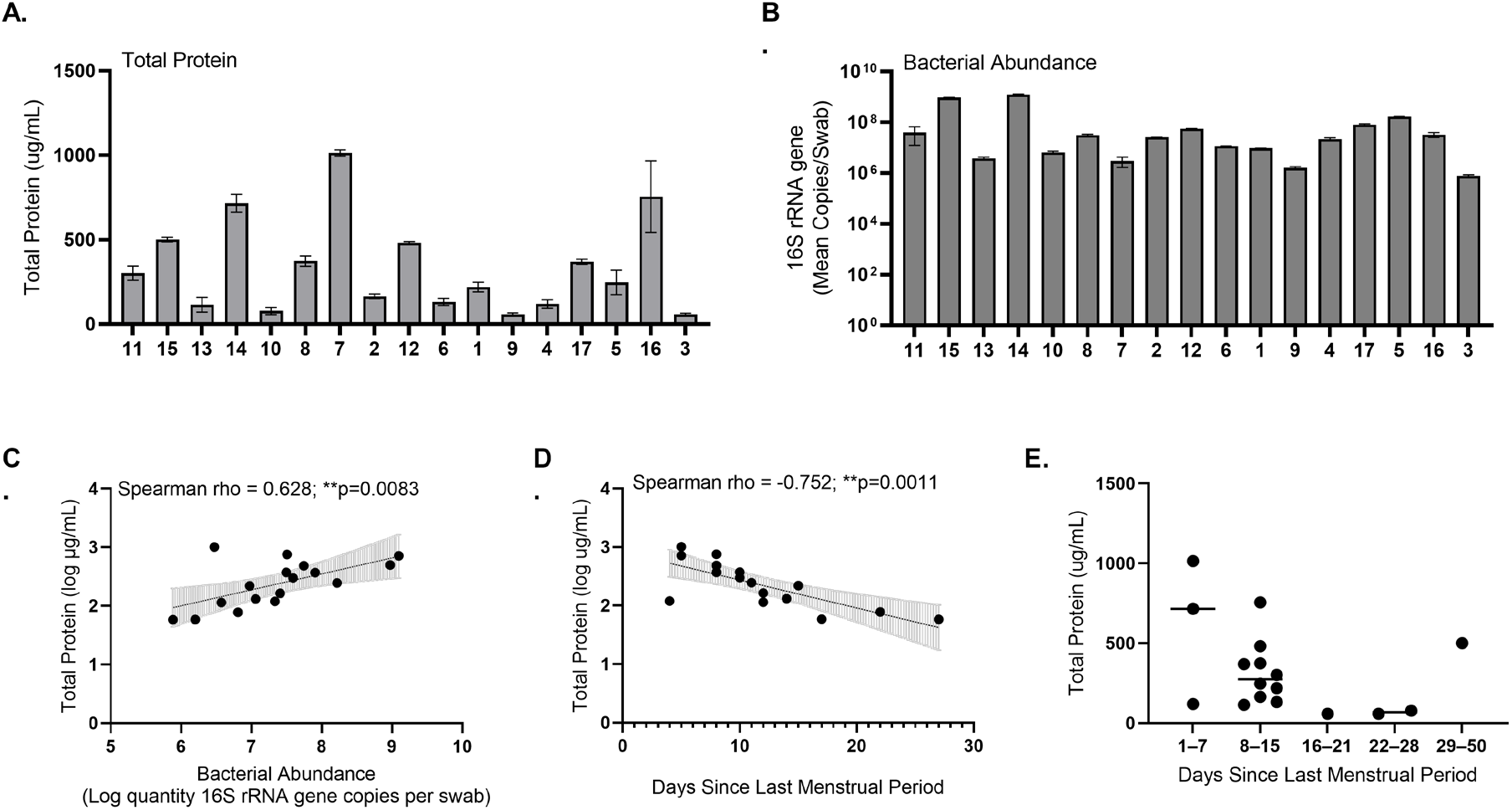
Correlates of total protein in vaginal swab supernatants. **(A)** Total protein concentration (µg/mL) measured in vaginal swab supernatants and presented as mean ± range from one independent experiment run in technical duplicate. **(B)** Bacterial abundance in vaginal swab cell pellets, as determined by broad-range 16S rRNA gene qPCR run in technical duplicate and presented as mean 16S rRNA gene copies per swab ± coefficient of variance. **(C–D)** Scatterplots of total protein concentration (µg/mL) as a function of **(C)** bacterial abundance (mean 16S rRNA gene copies/swab) and **(D)** days since last menstrual period (excluding one sample with >35 days). Regression lines were fitted using a generazlied linear model and error bars indicate 95% confidence intervals. Relationships were assessed with Spearman’s correlation, yielding the rho coefficients and p-values above each plot. **(E)** Distribution and mean of total protein concentration (µg/mL) for each swab supernatant sample by the week of the menstrual cycle in which the sample was collected (days since last menstrual period: 1–7 days, 8–15 days, 16–21 days, 22–28 days, or 29–50 days). Significance levels: ns=not statistically significant, *p≤0.05, **p≤0.01, ***p≤0.001, ****p≤0.0001.

**Figure S3.**
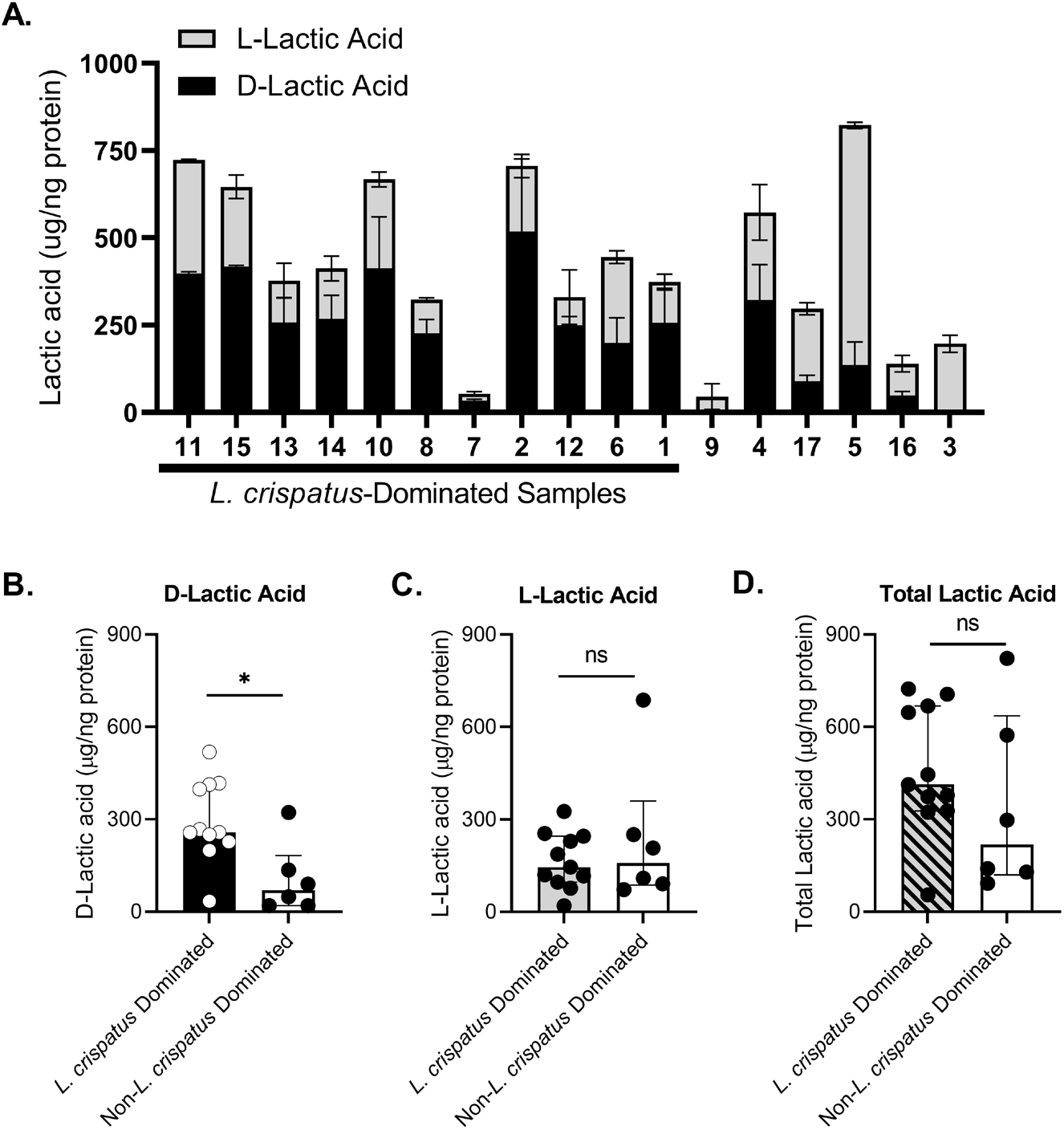
Elevated D-lactic acid levels in vaginal samples dominated by *L. crispatus*. **(A)** D-lactic acid, L-lactic acid and additive total lactic acid levels were quantified in vaginal swab supernatants (N=17) and normalized to total protein concentration. Results are presented as mean ± range of lactic acid from one independent experiment run in technical duplicate. **(B–D)** Distribution of lactic acid levels in *L. crispatus*-dominated (N=11) and non-*L. crispatus-*dominated (N=6) vaginal samples for **(B)** D-lactic acid **(C)** L-lactic acid and **(D)** total lactic acid presented as mean ± standard error of the mean. Statistical significance was assessed with a Mann-Whitney U-test: D-lactic acid *p=0.0202, L-lactic acid ns p>0.999. total lactic acid ns p=0.301. Significance levels: ns=not statistically significant, *p≤0.05, **p≤0.01, ***p≤0.001, ****p≤0.0001.

**Figure S4.**
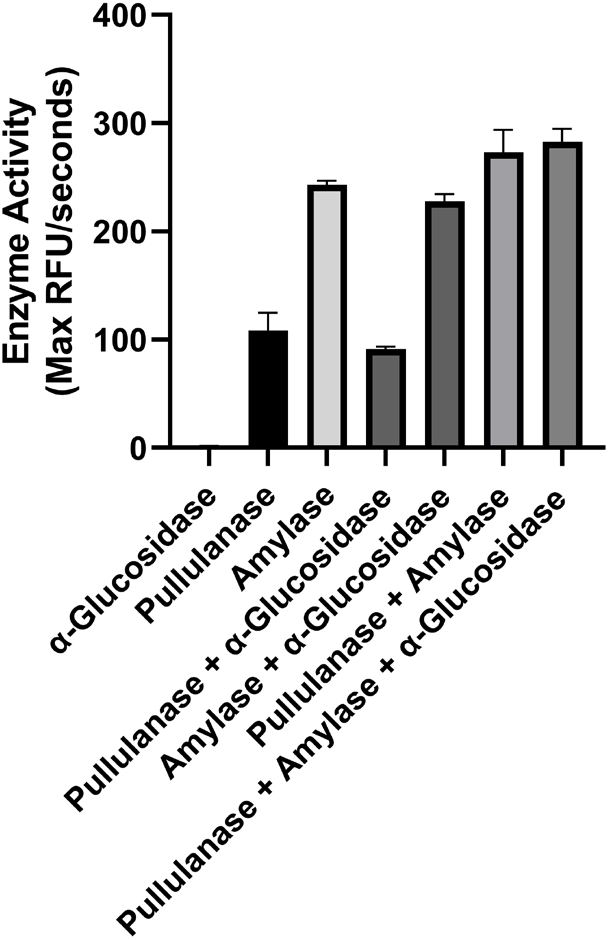
Enzyme activities detected in a fluorescent starch degradation assay. Fluorophore-conjugated starch was incubated with commercially available enzymes, α-glucosidase, pullulanase and α-amylase, alone and in combination. Degradation of the fluorescent starch substrate was quantified by measuring the increase in fluorescence (excitation/emission 485/527 nm) over a two-hour time course with reads every three minutes. The enzyme activity is expressed as the maximum relative fluorescence units (RFU)/seconds and presented as mean ± range from one independent experiment performed in technical duplicate.

**Figure S5.**
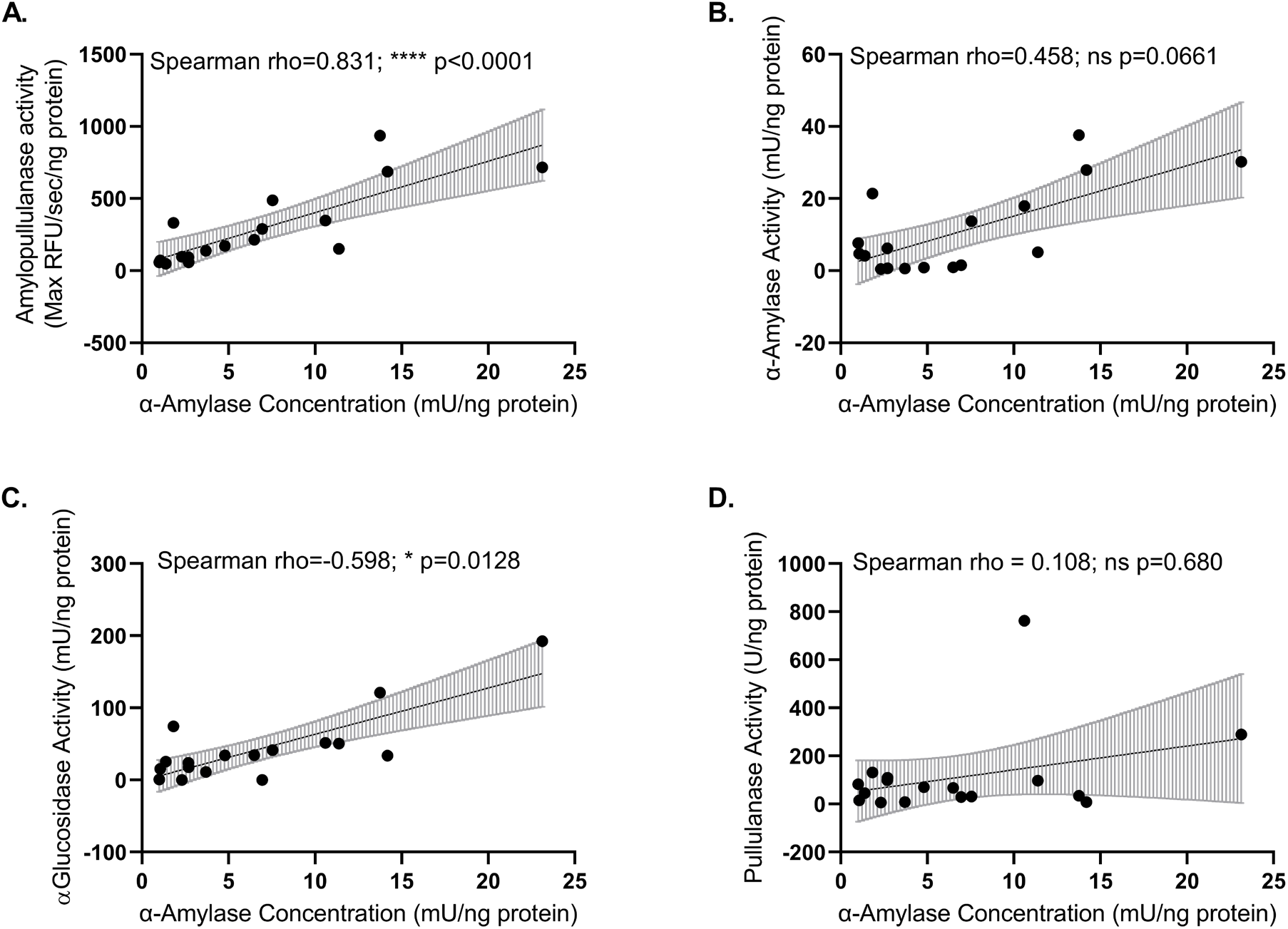
Human α-amylase correlates with amylopullulanase and α-glucosidase activities in vaginal swab supernatants, but not pullulanase activity. **(A–D)** Scatterplots of **(A)** amylopullulanase activity (max RFU/sec/ng protein) **(B)** α-amylase enzyme activity (mU/ng protein) **(C)** α-glucosidase enzyme activity (mU/ng protein) and **(D)** pullulanase enzyme activity (U/ng protein) as a function of human α-amylase enzyme concentration. All relationships were assessed with Spearman’s correlation, with the rho coefficient and p-value indicated above each scatterplot. Significance levels: ns=not statistically significant, *p≤0.05, **p≤0.01, ***p≤0.001, ****p≤0.0001.

**Figure S6.**
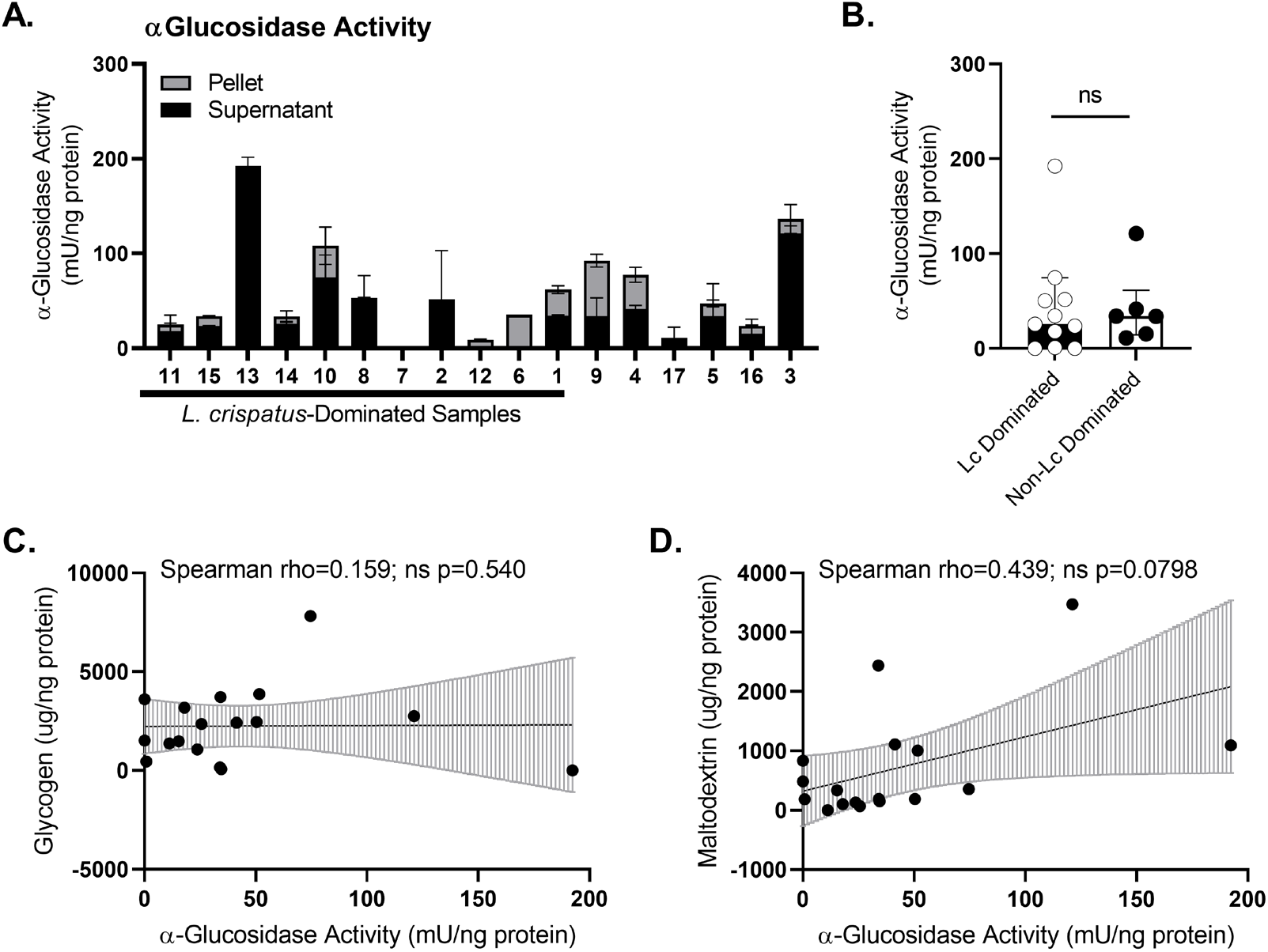
α-glucosidase activity correlates with maltodextrin levels in vaginal swab supernatants. **(A)** α-glucosidase enzyme activity was measured in vaginal swab supernatants (N=17) and in select vaginal swab cell pellets (all except samples 2, 7, 13 and 17; N=13). Results were normalized to total protein concentration and presented as mean mU/ng protein ± range from one independent experiment run in technical duplicate. **(B)** Distribution of α-glucosidase activity in *L. crispatus*-dominated (*Lc* Dominated; N=11) and non-*L. crispatus* dominated (non-*Lc* Dominated; N=6) vaginal supernatant samples presented as median ± interquartile range. Statistical significance between *Lc* Dominated and non-*Lc* Dominated samples was assessed with a Mann Whitney U-test (ns p=0.884). **(C–D)** Scatterplots of **(C)** glycogen and **(D)** maltodextrin levels as a function of α-glucosidase enzyme activity. Regression lines were fitted using a generalized linear model and error bars indicate 95% confidence intervals. Relationships were assessed with Spearman’s correlation, with the rho coefficient and p-value indicated above each scatterplot. Significance levels: ns=not statistically significant, *p≤0.05, **p≤0.01, ***p≤0.001, ****p≤0.0001.

**Figure S7.**
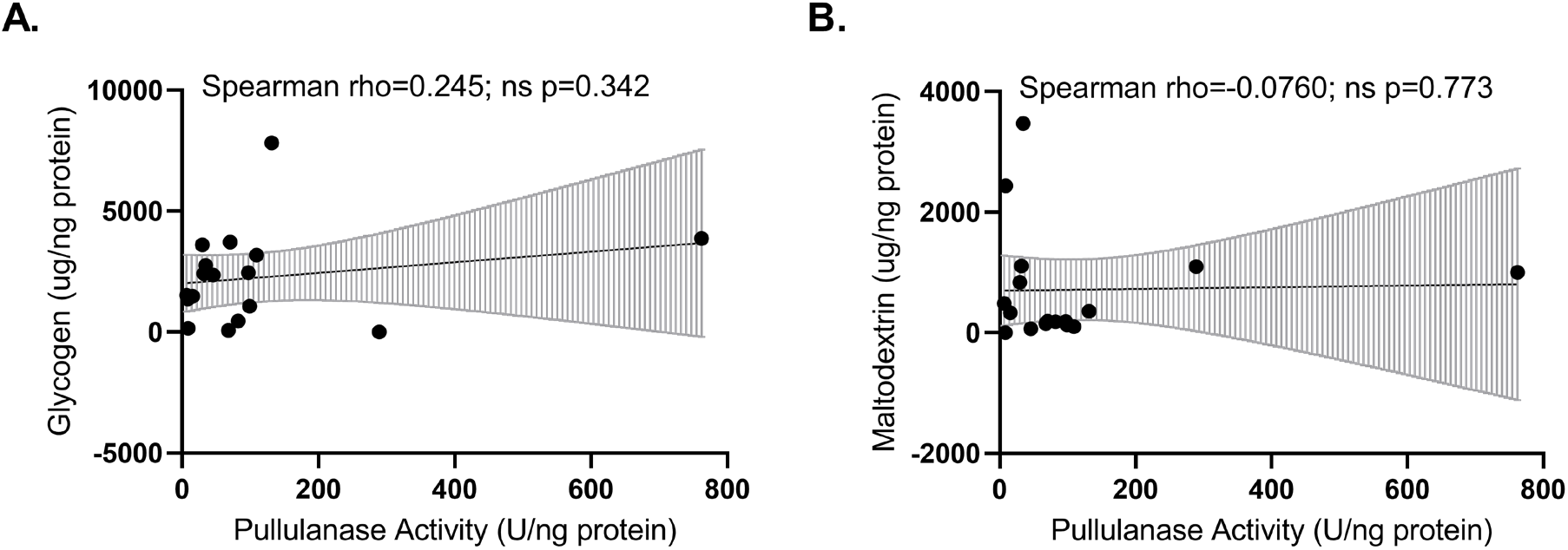
Pullulanase activity does not correlate with glycogen or maltodextrin levels in vaginal swab supernatants. Scatterplots of **(A)** glycogen or **(B)** maltodextrin measurements as a function of pullulanase activity. Regression lines were fitted using a generalized linear model with error bars indicating 95% confidence intervals. Relationships were assessed with Spearman’s correlation, with the rho coefficient and p-value indicated above each scatterplot. Significance levels: ns=not statistically significant, *p≤0.05, **p≤0.01, ***p≤0.001, ****p≤0.0001.

**Figure S8.**
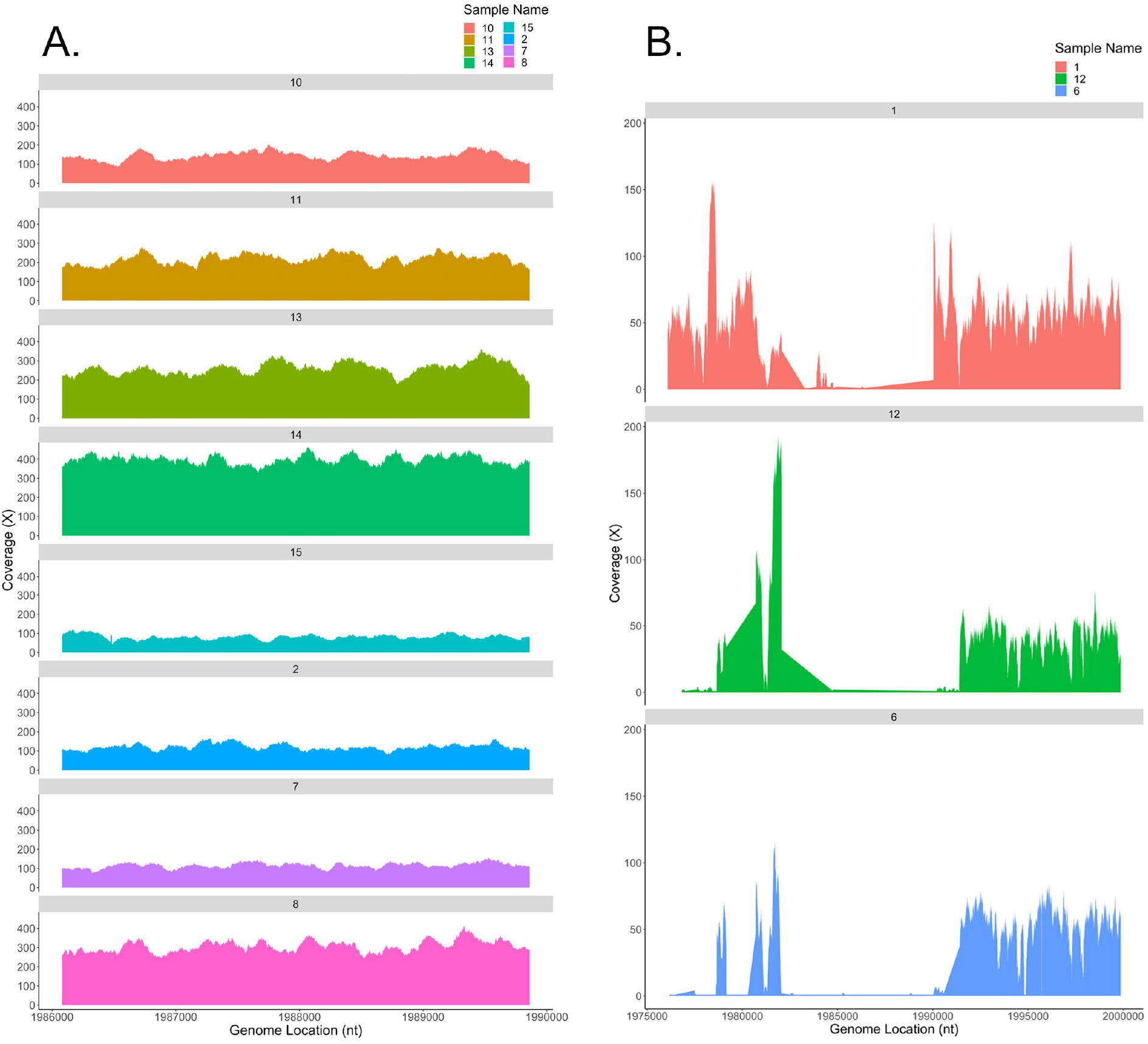
Coverage of the *L. crispatus pulA* gene and flanking regions in vaginal metagenomes dominated by *L. crispatus*. To further assess evidence of *pulA* gene presence/absence in vaginal metagenomes, we undertook read coverage analysis. Using unassembled reads, this approach served to confirm that our failure to detect *pulA* genes in some samples by BLASTn was due to low abundance/coverage, as opposed to assembly errors. **(A)** Depth of read coverage across the *pulA* gene (positions 1986082 to 1989861) when reads from each *L. crispatus-*dominated sample that produced a *pulA* BLASTn hit were mapped to the closed *L. crispatus* reference genome FFDAARGOS_743. Each sample that produced a *pulA* BLASTn hit displayed >100X coverage. **(B)** Read coverage of samples 1, 6 and 12, which all failed to produce a *pulA* BLASTn hit despite being dominated by *L. crispatus*, across a larger region consisting of 10,000 bases upstream and downstream of the *pulA* gene. High coverage regions, representative of what was seen across the broader *L. crispatus* reference genome (Tables S19–S20, Additional File 3), flanked a large low coverage area in the center of each graph that encompassed the *pulA* gene locus. This suggests a large genomic fragment that includes the *pulA* gene is missing in each of these genomes.

**Figure S9.**
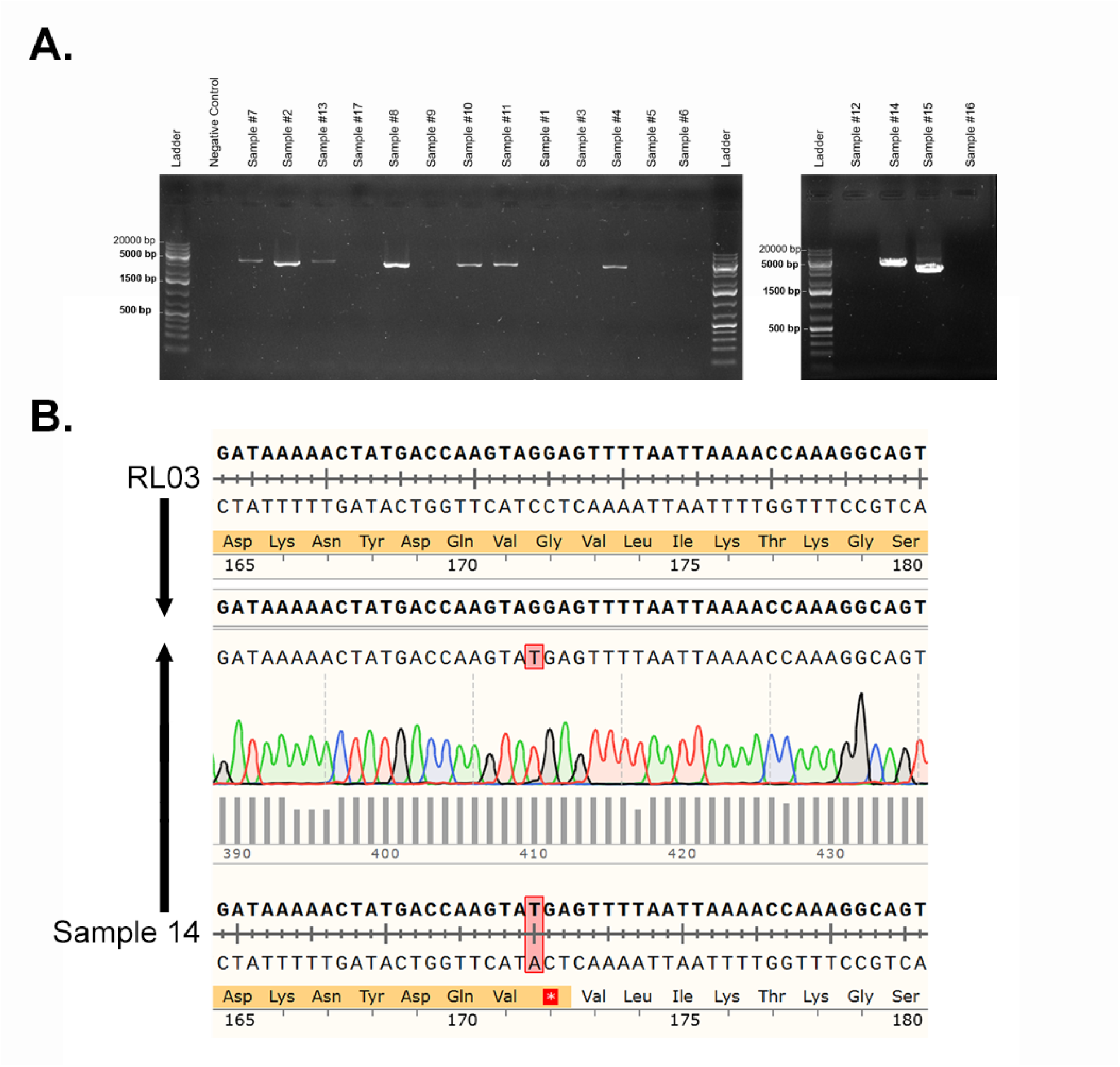
*L. crispatus pulA* gene amplification from vaginal swab samples. **(A)** Agarose gel (0.7%) electrophoresis of PCR products amplified using primers targeting nucleotides 74 to 3651 of the *L. crispatus* RL03 *pulA* gene (NCBI Accession: NZ_NKLQ01000285) with an expected product of 3578 bp. A ‘no DNA template’ negative control was included, as shown in lane #2 on the gel to the left. **(B)** Sanger sequencing traces generated by sequencing the *pulA* PCR product from Sample #14 from the 5’ end (primer binding at ∼74 bp) and aligning to the RL03 reference. The G514T single nucleotide polymorphism observed in this sample’s metagenomic assembly was evident, resulting in a premature stop codon at amino acid 172. Significance levels: ns=not statistically significant, *p≤0.05, **p≤0.01, ***p≤0.001, ****p≤0.0001.

**Figure S10.**
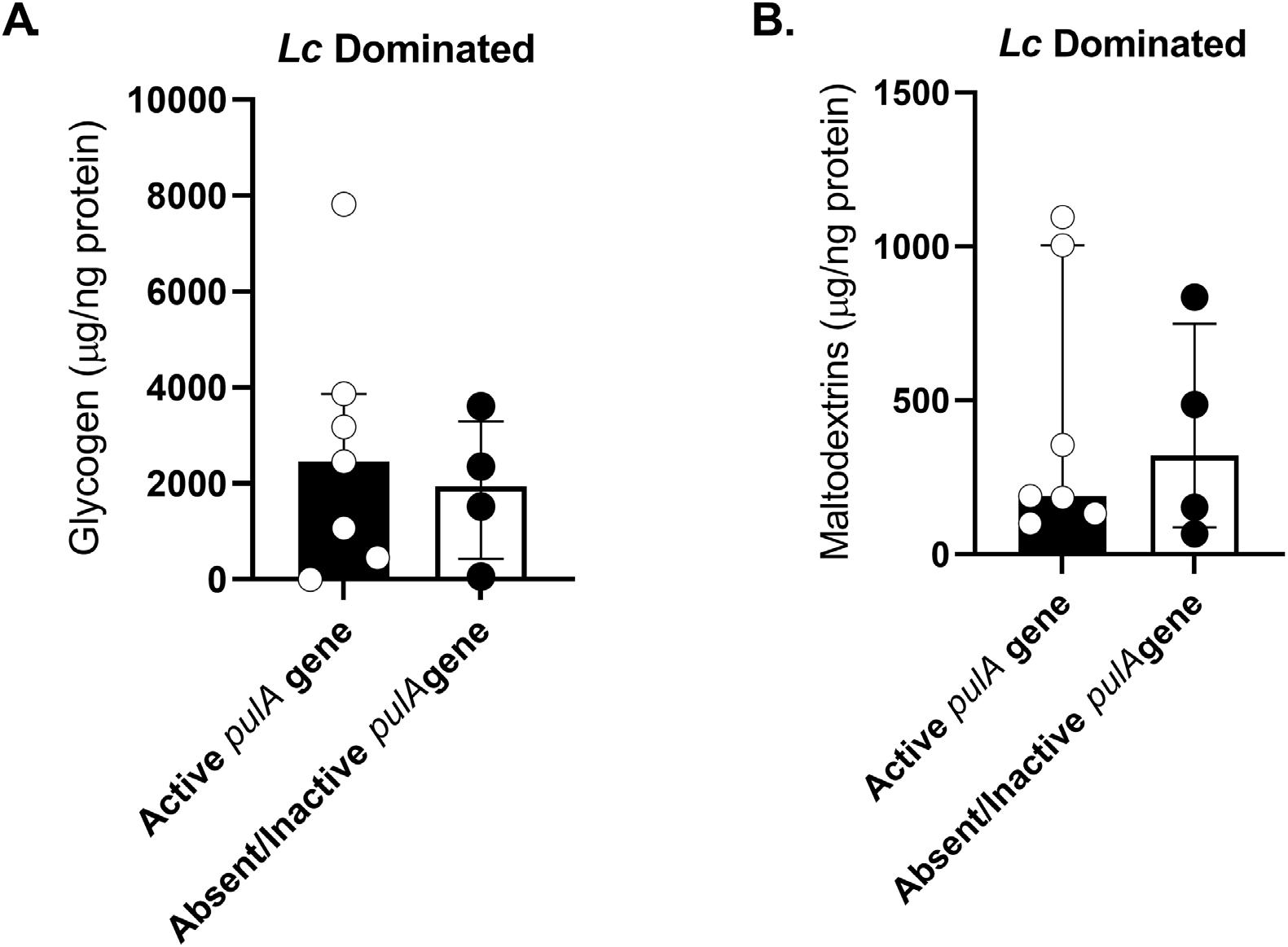
Glycogen and maltodextrin levels do not depend on whether *L. crispatus* encodes a functional pullulanase. Distribution of **(A)** glycogen (µg/ng protein) and **(B)** maltodextrin (µg/ng protein) levels in *L. crispatus*-dominated samples with a full-length *pulA* gene predicted to encode an active protein (N=7) and those missing or encoding a mutated *pulA* gene predicted to be inactivated (N=4). Data presented as median ± interquartile range and statistical significance between *Lc* Dominated samples with active vs. absent/inactive *pulA* was assessed with a Mann-Whitney U-test (glycogen ns p=0.788, maltodextrin ns p=0.788).

**Figure S11.**
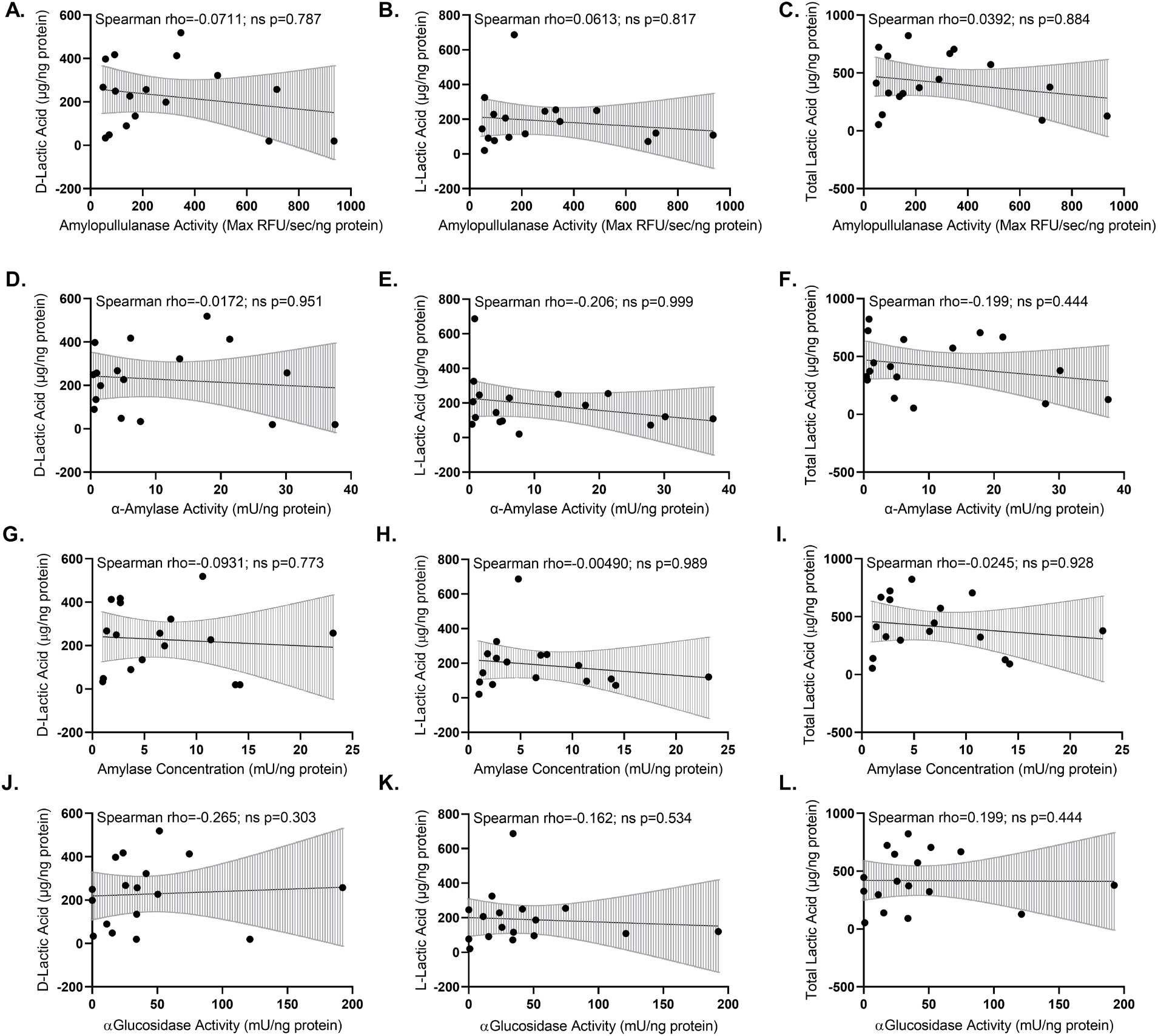
Amylase and α-glucosidase activities do not correlate with vaginal lactic acid. Scatterplots of D-lactic acid, L-lactic acid and total lactic acid measurements as a function of **(A–C)** amylopullulanase activity (Max RFU/sec/ng protein), **(D–F)** α-amylase activity (mU/ng protein**), (G–I)** human α-amylase concentration (mU/ng protein) or **(J–L)** α-glucosidase activity (mU/ng protein). Regression lines were fitted using a generalized linear model with error bars indicating 95% confidence intervals. Relationships were assessed with Spearman’s correlation, with the rho coefficient and p-value indicated above each scatterplot. Significance levels: ns=not statistically significant, *p≤0.05, **p≤0.01, ***p≤0.001, ****p≤0.0001.

**Figure S12.**
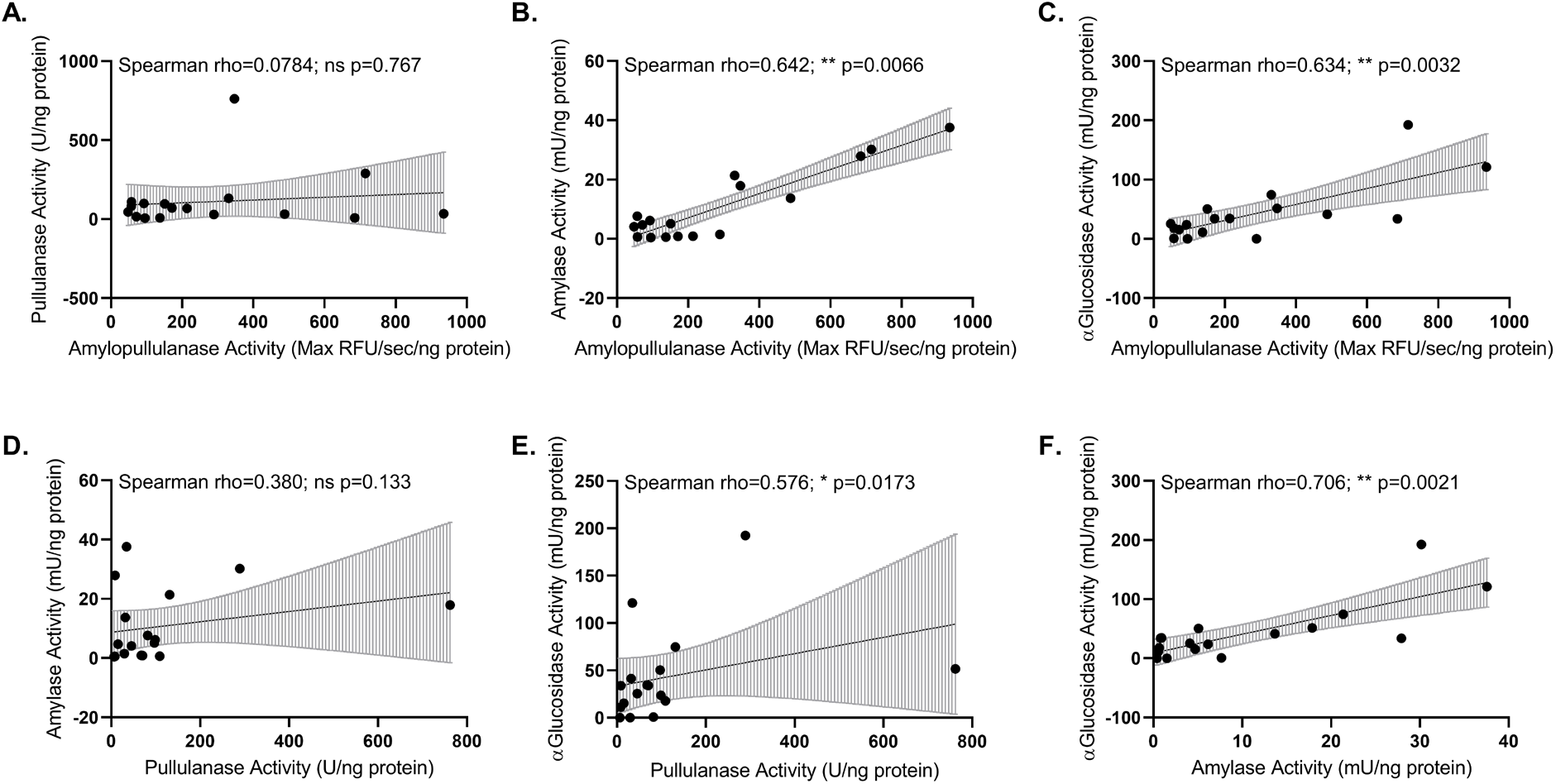
Correlations of enzyme activity in vaginal swab supernatants. **(A–C)** Scatterplots of **(A)** pullulanase activity (U/ng protein), **(B)** α-amylase activity (mU/ng protein) or **(C)** α-glucosidase activity (mU/ng protein) as a function of amylopullulanase activity (Max RFU/sec/ng protein). **(D,E)** Scatterplots of **(D)** α-amylase activity (mU/ng protein) or **(E)** α-glucosidase activity (mU/ng protein) as a function of pullulanase activity (U/ng protein) or **(F)** α-glucosidase activity (mU/ng protein) as a function of α-amylase activity (mU/ng protein). Regression lines were fitted using a generalized linear model with error bars indicating 95% confidence intervals. Relationships were assessed with Spearman’s correlation, with the rho coefficient and p-value indicated above each scatterplot. Significance levels: ns=not statistically significant, *p≤0.05, **p≤0.01, ***p≤0.001, ****p≤0.0001.

## References

1. Witkin, S. and I. Linhares, Why do lactobacilli dominate the human vaginal microbiota? BJOG: An International Journal of Obstetrics & Gynaecology, 2017. 124(4): p. 606–611.

2. Petrova, M.I., et al., Lactobacillus species as biomarkers and agents that can promote various aspects of vaginal health. Front Physiol, 2015. 6: p. 81.

3. Ravel, J., et al., Vaginal microbiome of reproductive-age women. Proceedings of the National Academy of Sciences, 2011. 108(Supplement 1): p. 4680–4687.

4. Ravel, J., et al., Vaginal microbiome of reproductive-age women. Proc Natl Acad Sci U S A, 2011. 108 Suppl 1: p. 4680–7.

5. Onderdonk, A.B., M.L. Delaney, and R.N. Fichorova, The Human Microbiome during Bacterial Vaginosis. Clin Microbiol Rev, 2016. 29(2): p. 223–38.

6. Hillier, S.L., et al., Association between bacterial vaginosis and preterm delivery of a low-birth-weight infant. The Vaginal Infections and Prematurity Study Group. N Engl J Med, 1995. 333(26): p. 1737–42.

7. Fettweis, J.M., et al., The vaginal microbiome and preterm birth. Nat Med, 2019. 25(6): p. 1012–1021.

8. Elovitz, M.A., et al., Cervicovaginal microbiota and local immune response modulate the risk of spontaneous preterm delivery. Nat Commun, 2019. 10(1): p. 1305.

9. Norenhag, J., et al., The vaginal microbiota, human papillomavirus and cervical dysplasia: a systematic review and network meta-analysis. BJOG, 2020. 127(2): p. 171–180.

10. Klein, C., et al., How the Cervical Microbiota Contributes to Cervical Cancer Risk in Sub-Saharan Africa. Front Cell Infect Microbiol, 2020. 10: p. 23.

11. Laniewski, P., et al., Linking cervicovaginal immune signatures, HPV and microbiota composition in cervical carcinogenesis in non-Hispanic and Hispanic women. Sci Rep, 2018. 8(1): p. 7593.

12. Sewankambo, N., et al., HIV-1 infection associated with abnormal vaginal flora morphology and bacterial vaginosis. Lancet, 1997. 350(9077): p. 546–50.

13. Wiesenfeld, H.C., et al., Bacterial vaginosis is a strong predictor of Neisseria gonorrhoeae and Chlamydia trachomatis infection. Clin Infect Dis, 2003. 36(5): p. 663–8.

14. Zhou, X., et al., Differences in the composition of vaginal microbial communities found in healthy Caucasian and black women. The ISME journal, 2007. 1(2): p. 121–133.

15. Zhou, X., et al., The vaginal bacterial communities of Japanese women resemble those of women in other racial groups. FEMS Immunology & Medical Microbiology, 2010. 58(2): p. 169–181.

16. Koumans, E.H., et al., The Prevalence of Bacterial Vaginosis in the United States, 2001–2004; Associations With Symptoms, Sexual Behaviors, and Reproductive Health. Sexually Transmitted Diseases, 2007. 34(11): p. 864–869.

17. Ness, R.B., et al., Can known risk factors explain racial differences in the occurrence of bacterial vaginosis? Journal of the National Medical Association, 2003. 95(3): p. 201.

18. Jespers, V., et al., The significance of Lactobacillus crispatus and L. vaginalis for vaginal health and the negative effect of recent sex: a cross-sectional descriptive study across groups of African women. BMC Infect Dis, 2015. 15: p. 115.

19. Royce, R.A., et al., Race/Ethnicity, Vaginal Flora Patterns, and pH During Pregnancy. Sexually Transmitted Diseases, 1999. 26(2): p. 96–102.

20. Benning, L., et al., Comparison of lower genital tract microbiota in HIV-infected and uninfected women from Rwanda and the US. PLoS One, 2014. 9(5): p. e96844.

21. Anukam, K.C., et al., Lactobacillus vaginal microbiota of women attending a reproductive health care service in Benin city, Nigeria. Sexually transmitted diseases, 2006. 33(1): p. 59–62.

22. Lennard, K., et al., Microbial Composition Predicts Genital Tract Inflammation and Persistent Bacterial Vaginosis in South African Adolescent Females. Infect Immun, 2018. 86(1).

23. Gosmann, C., et al., Lactobacillus-deficient cervicovaginal bacterial communities are associated with increased HIV acquisition in young South African women. Immunity, 2017. 46(1): p. 29–37.

24. Smart, S., A. Singal, and A. Mindel, Social and sexual risk factors for bacterial vaginosis. Sexually Transmitted Infections, 2004. 80(1): p. 58–62.

25. Fethers, K., et al., Bacterial vaginosis (BV) candidate bacteria: associations with BV and behavioural practices in sexually-experienced and inexperienced women. PloS one, 2012. 7(2): p. e30633.

26. Fethers, K.A., et al., Sexual risk factors and bacterial vaginosis: a systematic review and meta-analysis. Clinical Infectious Diseases, 2008. 47(11): p. 1426–1435.

27. Francis, S.C., et al., Results from a cross-sectional sexual and reproductive health study among school girls in Tanzania: high prevalence of bacterial vaginosis. Sexually transmitted infections, 2019. 95(3): p. 219–227.

28. Francis, S.C., et al., The Vaginal Microbiota Among Adolescent Girls in Tanzania Around the Time of Sexual Debut. Front Cell Infect Microbiol, 2020. 10: p. 305.

29. Mehta, S.D., et al., High Prevalence of Lactobacillus crispatus Dominated Vaginal Microbiome Among Kenyan Secondary School Girls: Negative Effects of Poor Quality Menstrual Hygiene Management and Sexual Activity. Front Cell Infect Microbiol, 2021. 11: p. 716537.

30. Atassi, F., et al., Lactobacillus strains isolated from the vaginal microbiota of healthy women inhibit Prevotella bivia and Gardnerella vaginalis in coculture and cell culture. FEMS Immunol Med Microbiol, 2006. 48(3): p. 424–32.

31. O’Hanlon, D.E., R.A. Come, and T.R. Moench, Vaginal pH measured in vivo: lactobacilli determine pH and lactic acid concentration. BMC Microbiol, 2019. 19(1): p. 13.

32. O’Hanlon, D.E., T.R. Moench, and R.A. Cone, Vaginal pH and microbicidal lactic acid when lactobacilli dominate the microbiota. PLoS One, 2013. 8(11): p. e80074.

33. O’Hanlon, D.E., T.R. Moench, and R.A. Cone, In vaginal fluid, bacteria associated with bacterial vaginosis can be suppressed with lactic acid but not hydrogen peroxide. BMC Infect Dis, 2011. 11: p. 200.

34. Aldunate, M., et al., Vaginal concentrations of lactic acid potently inactivate HIV. J Antimicrob Chemother, 2013. 68(9): p. 2015–25.

35. Gong, Z., et al., Lactobacilli inactivate Chlamydia trachomatis through lactic acid but not H2O2. PLoS One, 2014. 9(9): p. e107758.

36. Graver, M.A. and J.J. Wade, The role of acidification in the inhibition of Neisseria gonorrhoeae by vaginal lactobacilli during anaerobic growth. Ann Clin Microbiol Antimicrob, 2011. 10: p. 8.

37. Boskey, E.R., et al., Origins of vaginal acidity: high D/L lactate ratio is consistent with bacteria being the primary source. Hum Reprod, 2001. 16(9): p. 1809–13.

38. Witkin, S.S., et al., Influence of vaginal bacteria and D-and L-lactic acid isomers on vaginal extracellular matrix metalloproteinase inducer: implications for protection against upper genital tract infections. mBio, 2013. 4(4).

39. Mendes-Soares, H., et al., Comparative functional genomics of Lactobacillus spp. reveals possible mechanisms for specialization of vaginal lactobacilli to their environment. J Bacteriol, 2014. 196(7): p. 1458–70.

40. Tanizawa, Y., et al., Lactobacillus paragasseri sp. nov., a sister taxon of Lactobacillus gasseri, based on whole-genome sequence analyses. Int J Syst Evol Microbiol, 2018. 68(11): p. 3512–3517.

41. Mirmonsef, P., et al., Free glycogen in vaginal fluids is associated with Lactobacillus colonization and low vaginal pH. PLoS One, 2014. 9(7): p. e102467.

42. Mirmonsef, P., et al., Exploratory comparison of vaginal glycogen and Lactobacillus levels in premenopausal and postmenopausal women. Menopause, 2015. 22(7): p. 702–9.

43. Roach, P.J., et al., Glycogen and its metabolism: some new developments and old themes. Biochem J, 2012. 441(3): p. 763–87.

44. Spear, G.T., et al., Human alpha-amylase present in lower-genital-tract mucosal fluid processes glycogen to support vaginal colonization by Lactobacillus. J Infect Dis, 2014. 210(7): p. 1019–28.

45. Macklaim, J.M., et al., At the crossroads of vaginal health and disease, the genome sequence of Lactobacillus iners AB-1. Proc Natl Acad Sci U S A, 2011. 108 Suppl 1: p. 4688–95.

46. van der Veer, C., et al., Comparative genomics of human Lactobacillus crispatus isolates reveals genes for glycosylation and glycogen degradation: implications for in vivo dominance of the vaginal microbiota. Microbiome, 2019. 7(1): p. 49.

47. Wylie, J.G. and A. Henderson, Identity and glycogen-fermenting ability of lactobacilli isolated from the vagina of pregnant women. J Med Microbiol, 1969. 2(3): p. 363–6.

48. Stewart-Tull, D.E., Evidence That Vaginal Lactobacilli Do Not Ferment Glycogen. Am J Obstet Gynecol, 1964. 88: p. 676–9.

49. Martín, R., et al., Characterization of indigenous vaginal lactobacilli from healthy women as probiotic candidates. Int Microbiol, 2008. 11(4): p. 261–6.

50. Macklaim, J.M., et al., Comparative meta-RNA-seq of the vaginal microbiota and differential expression by Lactobacillus iners in health and dysbiosis. Microbiome, 2013. 1(1): p. 12.

51. Nunn, K.L., et al., Amylases in the Human Vagina. mSphere, 2020. 5(6).

52. Ma, B., et al., A comprehensive non-redundant gene catalog reveals extensive within-community intraspecies diversity in the human vagina. Nat Commun, 2020. 11(1): p. 940.

53. Bhandari, P., et al., Characterization of an alpha-Glucosidase Enzyme Conserved in Gardnerella spp. Isolated from the Human Vaginal Microbiome. J Bacteriol, 2021. 203(17): p. e0021321.

54. Yuh, T., et al., Sexually Transmitted Infections Among Kenyan Adolescent Girls and Young Women With Limited Sexual Experience. Front Public Health, 2020. 8: p. 303.

55. Khot, P.D., et al., Development and optimization of quantitative PCR for the diagnosis of invasive aspergillosis with bronchoalveolar lavage fluid. BMC Infect Dis, 2008. 8: p. 73.

56. Martin, M., Cutadapt removes adapter sequences from high-throughput sequencing reads. EMBnet.journal, 2011. 17(1): p. 10–12.

57. Schmieder, R. and R. Edwards, Quality control and preprocessing of metagenomic datasets. Bioinformatics, 2011. 27(6): p. 863–4.

58. Rotmistrovsky, K.a.A., R., BMTagger: Best Match Tagger for removing human reads from metagenomics datasets. ftp://ftp.ncbi.nlm.nih.gov/pub/agarwala/bmtagger/.

59. Beghini, F., et al., Integrating taxonomic, functional, and strain-level profiling of diverse microbial communities with bioBakery 3. Elife, 2021. 10.

60. Wickham, H., ggplot2: Elegant Graphics for Data Analysis. 2016, Springer-Verlag New York.

61. A language and environment for statistical computing. R Foundation for Statistical Computing, Vienna, Austria. URL https://www.R-project.org/. R Core Team, 2017.

62. Oksanen, J. 2020; Available from: https://cran.r-project.org/package=vegan.

63. Kolde, R. pheatmap: Pretty Heatmaps. 2019; Available from: https://cran.r-project.org/package=pheatmap.

64. Neuwirth, E. RColorBrewer: ColorBrewer Palettes. 2014; Available from: https://cran.r-project.org/package=RColorBrewer.

65. Uritskiy, G.V., J. DiRuggiero, and J. Taylor, MetaWRAP-a flexible pipeline for genome-resolved metagenomic data analysis. Microbiome, 2018. 6(1): p. 158.

66. Bankevich, A., et al., SPAdes: a new genome assembly algorithm and its applications to single-cell sequencing. J Comput Biol, 2012. 19(5): p. 455–77.

67. Gurevich, A., et al., QUAST: quality assessment tool for genome assemblies. Bioinformatics, 2013. 29(8): p. 1072–5.

68. Kang, D.D., et al., MetaBAT, an efficient tool for accurately reconstructing single genomes from complex microbial communities. PeerJ, 2015. 3: p. e1165.

69. Wu, Y.W., B.A. Simmons, and S.W. Singer, MaxBin 2.0: an automated binning algorithm to recover genomes from multiple metagenomic datasets. Bioinformatics, 2016. 32(4): p. 605–7.

70. Sharon, I., et al., Time series community genomics analysis reveals rapid shifts in bacterial species, strains, and phage during infant gut colonization. Genome Res, 2013. 23(1): p. 111–20.

71. Parks, D.H., et al., CheckM: assessing the quality of microbial genomes recovered from isolates, single cells, and metagenomes. Genome Res, 2015. 25(7): p. 1043–55.

72. Seemann, T., Prokka: rapid prokaryotic genome annotation. Bioinformatics, 2014. 30(14): p. 2068–9.

73. Chaumeil, P.A., et al., GTDB-Tk: a toolkit to classify genomes with the Genome Taxonomy Database. Bioinformatics, 2019.

74. Ferretti, P., et al., Mother-to-Infant Microbial Transmission from Different Body Sites Shapes the Developing Infant Gut Microbiome. Cell Host Microbe, 2018. 24(1): p. 133–145 e5.

75. Pasolli, E., et al., Extensive Unexplored Human Microbiome Diversity Revealed by Over 150,000 Genomes from Metagenomes Spanning Age, Geography, and Lifestyle. Cell, 2019. 176(3): p. 649–662 e20.

76. Human Microbiome Project, C., A framework for human microbiome research. Nature, 2012. 486(7402): p. 215–21.

77. Altschul, S.F., et al., Basic local alignment search tool. J Mol Biol, 1990. 215(3): p. 403–10.

78. Camacho, C., et al., BLAST+: architecture and applications. BMC Bioinformatics, 2009. 10: p. 421.

79. Sichtig, H., et al., FDA-ARGOS is a database with public quality-controlled reference genomes for diagnostic use and regulatory science. Nat Commun, 2019. 10(1): p. 3313.

80. Langmead, B. and S.L. Salzberg, Fast gapped-read alignment with Bowtie 2. Nat Methods, 2012. 9(4): p. 357–9.

81. Li, H., et al., The Sequence Alignment/Map format and SAMtools. Bioinformatics, 2009. 25(16): p. 2078–9.

82. Seemann, T., Prokka: rapid prokaryotic genome annotation. Bioinformatics, 2014. 30(14): p. 2068–2069.

83. Edgar, R.C., MUSCLE: multiple sequence alignment with high accuracy and high throughput. Nucleic Acids Res, 2004. 32(5): p. 1792–7.

84. Stamatakis, A., RAxML version 8: a tool for phylogenetic analysis and post-analysis of large phylogenies. Bioinformatics, 2014. 30(9): p. 1312–1313.

85. Stamatakis, A. Phylogenetic models of rate heterogeneity: a high performance computing perspective. in Proceedings 20th IEEE International Parallel & Distributed Processing Symposium. 2006.

86. Felsenstein, J., Confidence Limits on Phylogenies: An Approach Using the Bootstrap. Evolution, 1985. 39(4): p. 783–791.

87. Klatt, N.R., et al., Vaginal bacteria modify HIV tenofovir microbicide efficacy in African women. Science, 2017. 356(6341): p. 938–945.

88. Nunn, K.L., et al., Vaginal Glycogen, Not Estradiol, Is Associated With Vaginal Bacterial Community Composition in Black Adolescent Women. J Adolesc Health, 2019. 65(1): p. 130–138.

89. Mirmonsef, P., et al., An exploratory comparison of vaginal glycogen and Lactobacillus levels in pre-and post-menopausal women. Menopause (New York, NY), 2015. 22(7): p. 702.

90. Mitchell, C.M., et al., Vaginal microbiota and genitourinary menopausal symptoms: a cross-sectional analysis. Menopause, 2017. 24(10): p. 1160–1166.

91. Xu, P., et al., Biotechnology and bioengineering of pullulanase: state of the art and perspectives. World J Microbiol Biotechnol, 2021. 37(3): p. 43.

92. Shim, J.H., et al., Role of maltogenic amylase and pullulanase in maltodextrin and glycogen metabolism of Bacillus subtilis 168. J Bacteriol, 2009. 191(15): p. 4835–44.

93. Nisha, M. and T. Satyanarayana, Recombinant bacterial amylopullulanases: developments and perspectives. Bioengineered, 2013. 4(6): p. 388–400.

94. Cruickshank, R. and A. Sharman, The biology of the vagina in the human subject. BJOG: An International Journal of Obstetrics & Gynaecology, 1934. 41(2): p. 208–226.

95. Hickey, R.J., et al., Vaginal microbiota of adolescent girls prior to the onset of menarche resemble those of reproductive-age women. MBio, 2015. 6(2).

96. Spear, G.T., et al., Effect of pH on Cleavage of Glycogen by Vaginal Enzymes. PLoS One, 2015. 10(7): p. e0132646.

97. Woolston, B.M., et al., Characterization of vaginal microbial enzymes identifies amylopullulanases that support growth of Lactobacillus crispatus on glycogen. bioRxiv, 2021.

98. Fontana, F., et al., Probiogenomics Analysis of 97 Lactobacillus crispatus Strains as a Tool for the Identification of Promising Next-Generation Probiotics. Microorganisms, 2020. 9(1).

99. Mitra, A., et al., Comparison of vaginal microbiota sampling techniques: cytobrush versus swab. Sci Rep, 2017. 7(1): p. 9802.

100. Mitchell, C.M., et al., Effect of sexual debut on vaginal microbiota in a cohort of young women. Obstet Gynecol, 2012. 120(6): p. 1306–13.

101. Langhaug, L.F., L. Sherr, and F.M. Cowan, How to improve the validity of sexual behaviour reporting: systematic review of questionnaire delivery modes in developing countries. Trop Med Int Health, 2010. 15(3): p. 362–81.

